# Spatiotemporal patterns of sleep spindle activity in human anterior thalamus and cortex

**DOI:** 10.1101/2022.03.25.485812

**Authors:** Hannah Bernhard, Frederic L. W. V. J. Schaper, Marcus L. F. Janssen, Erik D. Gommer, Bernadette M. Jansma, Vivianne Van Kranen-Mastenbroek, Rob P. W. Rouhl, Peter de Weerd, Joel Reithler, Mark J. Roberts, DBS study group

## Abstract

Sleep spindles (8 - 16 Hz) are transient electrophysiological events during non-rapid eye movement sleep. While sleep spindles are routinely observed in the cortex using scalp electroencephalography (EEG), recordings of their thalamic counterparts have not been widely studied in humans. Based on a few existing studies, it has been hypothesized that spindles occur as largely local phenomena. We investigated intra-thalamic and thalamocortical spindle co-occurrence, which may underlie thalamocortical communication. We obtained scalp EEG and thalamic recordings from 7 patients that received bilateral deep brain stimulation (DBS) electrodes to the anterior thalamus for the treatment of drug resistant focal epilepsy. Spindles were categorized into subtypes based on their main frequency (i.e., slow (10±2 Hz) or fast (14±2 Hz)) and their level of thalamic involvement (spanning one channel, or spreading uni- or bilaterally within the thalamus). For the first time, we contrasted observed spindle patterns with permuted data to estimate random spindle co-occurrence. We found that multichannel spindle patterns were systematically coordinated at the thalamic and thalamocortical level. Importantly, distinct topographical patterns of thalamocortical spindle overlap were associated with slow and fast subtypes of spindles. These observations provide further evidence for coordinated spindle activity in thalamocortical networks.

**Highlights:**

- Sleep spindles were measured in human anterior thalamus and on the scalp
- Both fast and slow spindles occurred in the anterior thalamus
- > 25% of spindles spanned multiple channels in thalamus and cortex
- A novel statistical approach confirmed that spindle co-occurrences were not random
- Cortical spindle patterns depended on thalamic involvement and spindle frequency

## Introduction

Sleep spindles are transient (0.3 – 2 s) electrophysiological patterns (8 – 16 Hz) and a characteristic of non-rapid eye movement (NREM) sleep in humans and other mammals (Fernandez & Lüthi, 2020). Recent research suggests that spindles may be involved in coordinating information transfer in hippocampal and neocortical networks to support memory consolidation and learning (Andrillon & Kouider, 2020; Klinzing et al., 2019; Latchoumane et al., 2017; Ngo et al., 2020; Staresina et al., 2015). Spindles across the brain can occur in a number of ways. They could be locally restricted to specific (cortical) sites or they could spread from thalamus to cortex in a set manner. Both options would preclude spindles from flexibly coordinating information transfer. Alternatively, spindles might follow function-specific spatiotemporal patterns. To the best of our knowledge, the extent to which spindles systematically co-occur within thalamic and across thalamocortical networks is unknown. Here, we investigated in humans whether sleep spindle activity in anterior thalamus and cortex follows coordinated spatiotemporal patterns.

To date, only few studies on human thalamic spindles exist. The reasons for this are twofold. Most human spindle research has relied on scalp electroencephalography (EEG), whose limited spatial resolution does not permit inferences about thalamic spindles. Thalamic signals can only be recorded using invasive procedures, but invasive thalamic recordings are generally performed during awake brain surgery or under anesthesia (Schaper et al., 2019). Deep brain recordings acquired via externalized deep brain stimulation (DBS) leads (Schreiner, Kaufmann, et al., 2021; Szalárdy et al., 2021), provide a safe (Feldmann et al., 2021) opportunity to study thalamic spindles in clinical populations of patients with epilepsy. Several studies have performed recordings from the posterior thalamus and cortex. For example, Mak-McCully et al. showed that thalamic spindles can precede spindles in neocortex (Mak-McCully et al., 2017). Bastuji et al. reported a tendency of localized thalamic spindles to also have more localized cortical counterpart, while diffuse thalamic spindles were correspondingly more diffusely represented in the cortex (Bastuji et al., 2020). Based on these data, the authors have inferred coordinated spindle activity across networks. However, their analysis did not account for random temporal overlap of spindle activity. In addition, these studies did not differentiate between so called slow (8 – 12 Hz) and fast (12 – 16 Hz) spindles. This differentiation seems relevant, because different types of spindles might have different functions. Fast spindles have been linked to memory processes (Ngo et al., 2013). The functional role as well as the physiology of slow spindles is understudied, as the majority of studies filtered in the spindle frequency range of 11 to 16 Hz (Bastuji et al., 2020; Mak-McCully et al., 2017). Furthermore, due to clinical constraints, these studies focused on the unilateral posterior thalamus. Consequently, the spindle behavior within the anterior thalamus and between anterior thalamus and cortex remains unknown. A characterization of spindles in the anterior thalamus is pertinent, because of its relation to memory function in animals (Aggleton et al., 2010; Frost et al., 2021) and humans (Sweeney-Reed et al., 2014, 2017, 2021).

Here, we addressed these issues and took a statistical approach to measure patterns of slow and fast spindle (co-)occurrence in the anterior thalamus and cortex by controlling for random spindle co-occurrence using permutation testing. This approach allowed us to compare temporal spindle coupling observed in the data with distributions of random spindle co-occurrence, thereby informing whether spindles in thalamocortical loops occur in systematically coordinated patterns. We recorded data in 7 individuals with epilepsy who underwent implantation of DBS quadripolar electrodes targeting the anterior nucleus of the thalamus (ANT). The setup of bilateral thalamic depth electrodes allowed us to assess co-occurrences of thalamic spindles within and across hemispheres. Simultaneous 19-channel scalp EEG recordings made it possible to study thalamocortical spindle co-occurrence globally across the cortex. We categorized spindles based on their main frequency as slow (10±2 Hz) or fast (14±2 Hz) spindles, as well as their level of thalamic involvement (spindles restricted to one channel, or spanning more than one channel unilaterally or bilaterally). We found that both slow and fast thalamic and thalamocortical spindles were more likely to co-occur across channels than expected by chance. Additionally, distinct topographical patterns of spindle overlap were associated with the different spindle subtypes. These observations provide support that slow and fast spindle activity are systematically coordinated in thalamocortical circuits.

## Materials & Methods

### Study design

The participants included in this study were admitted to Maastricht University Medical Centre for the implantation of DBS as treatment for drug-resistant focal epilepsy. In- and exclusion criteria followed the FDA and CE mark of the SANTE trial and are described elsewhere (Fisher et al., 2010; Salanova et al., 2015; Schaper et al., 2020). Patients were monitored with EEG and intracranial registration of their DBS electrodes for 22 hours after DBS implantation, including one night of sleep. The study procedure (DBSSEPI study (Dutch Trial Register NL4440 (NTR4562))) was approved by the medical ethical committee of Maastricht University Medical Centre, the Netherlands (ID: METC 14-4-126), and was conducted in accordance with the declaration of Helsinki.

### Participants

Seven individuals with drug-resistant focal epilepsy were included in this study (2 female; mean age 35 years; range 18 – 59 years). Six other individuals, who were part of the DBSSEPI study (Schaper et al., 2019, 2020) were excluded due to missing scalp electrodes and/or seizures during the night of recording.

### DBS surgery

As part of standard clinical practice, all participants had a pre-operative 3-T MRI (Philips, Eindhoven, The Netherlands) or 1.5-T MRI (Philips, Eindhoven, The Netherlands) in case of an implanted vagal nerve stimulator for stereotactic planning of the electrode trajectory. The sequences used were a 3D T1 with gadolinium, axial T2 and a T1 inversion recovery.

DBS electrode implantation was performed by directly targeting the ANT on pre-operative MRI. The most distal contacts were placed at the termination point of the mammillothalamic tract (MTT) in the ANT, also called the ANT-MTT junction (Lehtimäki et al., 2019). DBS surgery was performed under general anesthesia with remifentanil and propofol using a Leksell stereotactic frame, under guidance of intraoperative microelectrode recording. Details of the DBS surgery and lead placement are described elsewhere (Schaper et al., 2019, 2020).

### DBS lead localization

DBS leads were localized in a common space (Montreal Neurological Institute (MNI) 1522009b template (Fonov et al., 2011)) using Lead-DBS software (https://www.lead-dbs.org) with default settings (Horn et al., 2019), in line with previous studies (Horn et al., 2017). In short, pre-operative T1 / T2 MRI sequences and post-operative MRI / CT images (suppl. table 1) were linearly co-registered using SPM (Friston et al., 1994), version 12. Co-registration was further refined using the *brainshift correction* option (Schönecker et al., 2009) and images were normalized to MNI space using the Advanced Normalization Tool (http://stnava.github.io/ANTs/; Avants et al., 2008). DBS electrode trajectories and contacts were automatically pre-localized and manually refined using Lead-DBS. DBS lead trajectories were visualized in MNI space in relation to a thalamic mask, derived from the Thomas atlas (Su et al., 2019) and can be seen in figure 1.

**Figure 1.**
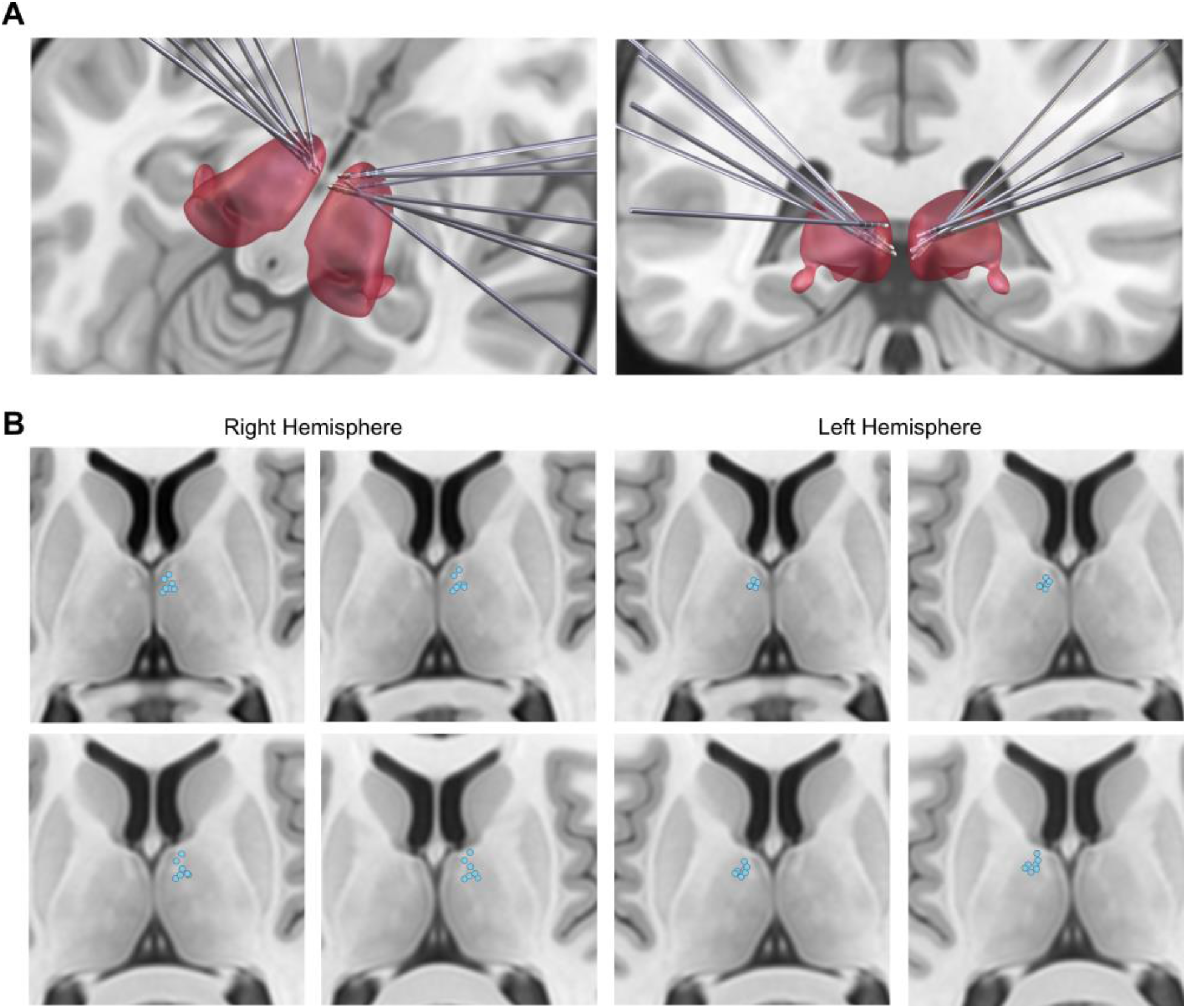
DBS lead locations and anterior thalamic recording sites in a common atlas space. **A)** Bilateral trajectories of implanted DBS leads for all participants (n=7) in transversal (left) and coronal (right) view. Note that each lead contained four electrode contacts. **B)** Contact positions (blue dots) of all participants projected onto a thalamic mask derived from the Thomas atlas (Su et al., 2019). MNI coordinates of each contact can be found in supplementary table 1.

### Data acquisition of electrophysiological recordings

Thalamic local field potentials (LFPs) were acquired using quadripolar externalized DBS leads (model 3389, Medtronic, Dublin, Ireland). Scalp EEG was measured using a 19-channel setup, with Ag/AgCl cup electrodes placed according to the standard 10-20 system. Data were recorded in Brain RT version 3.3 at a sampling rate of 2048 Hz, using Braintronics BrainBox-1166-64.

### Sleep scoring

Sleep scoring was performed in Brain RT version 3.3. For three out of seven participants, eye movement activity was measured with electro-oculogram (EOG) electrodes placed on the outer corners of the eyes, recording horizontal (HEOG) and vertical (VEOG) eye movements, respectively. For sleep scoring, a bipolar montage was applied to HEOG and VEOG channels, respectively. Four patients did not have EOG recordings. In this case, the frontal scalp EEG channels (Fp1 and Fp2, as well as F7 and F8, respectively) were re-referenced to bipolar derivations to show maximum eye movement activity for the polysomnographic scoring (i.e., Fp1 to F7, and Fp2 to F8), enabling the identification of wake and phasic REM periods. Channels F3, C3, O1, were re-referenced to the right mastoid, channels C4 and O2 to the left mastoid. Using a notch-filter at 50 Hz and a window size of 30s, a sleep scoring expert (H.B.) manually scored the data into sleep stages according to the American Academy of Sleep Medicine (AASM) criteria (Berry et al., 2015).

### Data Preprocessing

The Fieldtrip toolbox (Oostenveld, Fries, Maris, & Schoffelen, 2011; http://fieldtriptoolbox.org), Sleeptrip toolbox (), MATLAB R2018a and 2021b (Mathworks Inc., Sherbom, MA) were used for data analysis. Hypnograms and continuous data were exported from Brain RT to Matlab. A high pass filter at 0.4 Hz (Butterworth, two-pass filter, 5^th^ order) was applied to the continuous data, before down-sampling to 512 Hz. Electrodes were checked manually and, in case of continuous high noise levels, excluded. This was only the case for one electrode (A2) in one participant.

Scalp EEG activity was then re-referenced to the average of the mastoid electrodes (A1, A2), or to the average of all scalp electrodes for the patient missing A2. For the thalamic electrodes, bipolar derivations were computed between neighboring electrodes, resulting in 6 thalamic channels per participant (3 per hemisphere). We chose bipolar derivations to obtain a more localized signal. Where a certain thalamic nucleus is named in the supplementary material of this paper, one or both of the electrodes in the bipolar derivation were localized within this nucleus. However, given the length (1.5mm), diameter (1.27 mm) and spacing (0.55 mm) of electrode contacts, bipolar derivations may not be located in one nucleus but span several.

Both thalamic LFP and scalp EEG data were subjected to a low pass filter at 100Hz (Butterworth, two-pass, 5^th^ order), and a line noise filter at 50 Hz to reduce electrical noise. Subsequently, the continuous data were transformed into 30 s epochs and linked with the sleep hypnogram. Epochs with manually identified artifacts (e.g., visibly noisy signal) or arousals were excluded, as well as any epochs not pertaining to NREM sleep (N2 and N3, respectively).

### Spindle detection

Automatized spindle detection was performed in Sleeptrip, based on a pipeline used in previous literature (Weber et al., 2020). Spindles were detected in NREM sleep. To summarize, data from NREM segments were subjected to a FIR filter with a peak-frequency of 10 ± 2 Hz for slow and 14 ± 2 Hz for fast spindles, respectively. Next, the absolute value of the Hilbert transform of the filtered signal was taken to detect the spindles. Segments in which the envelope surpassed the threshold of the mean ±1.5 SD were marked as putative spindles. If the threshold crossings fell into the duration limits of 0.3 to 2 s, and a second threshold of 2.25 SD was passed at least once, these segments were considered as spindles. Spindles were not merged if they closely followed each other. Spindle detection was implemented separately for each channel and participant.

### Spindle detection control analysis

To verify that detected time windows included true spindles and not false positives, e.g. due to broadband frequency changes, we implemented a control analysis based on Ngo et al. (2020). This control analysis was done separately for slow and fast spindles. For each channel, time-frequency representations (TFRs) of each spindle were created using wavelets (5 cycles; 5 to 15 Hz for slow spindles; 9 to 19 Hz for fast spindles; in 0.5 Hz steps, time window of ± 0.75 s in 2 ms steps). Per event, frequencies were averaged between - 0.5 to + 0.5 s. Then, the *findpeaks* function was used on the resulting spectrogram. A putative spindle was considered a true spindle when the peak surpassed 0.2 prominence, and if the peak was between 8 to 12 Hz (for slow spindles) or 12 to 16 Hz (for fast spindles). All false positives were excluded from further analyses. For the number of detected spindle events and those that passed the control analysis, see table 2.

### Spindle characteristics

To quantify spindle parameters in the anterior thalamus, we computed several descriptive measures per channel and per participant for slow and fast spindles separately, namely: Spindle duration (difference of spindle offset and onset); spindle density (number of spindles per minute) and spindle amplitude (maximum peak to trough potential difference). Putative differences of spindle characteristics between fast and slow spindles were compared using paired t-tests per channel, setting the two-tailed significance level at α = 0.05. In a separate step, we examined whether channels in certain nuclei differed in spindle characteristics (supplementary material and supplementary figure 2), but caution against their overinterpretation due to several factors: Our definition of the exact nucleus can only be viewed as an approximation, the number of electrode contacts differ between thalamic nuclei, and the age range of the sample may lead to variability within a given nucleus.

To visualize the presence of slow and fast spindles in the thalamic LFP, we selected time-windows of ±1 s around the midpoint of algorithmically detected slow and fast spindles. Next, the TFR of each spindle was created using wavelets (5 cycles; 6 to 18 Hz, time window of ± 0.75 s in 1 ms steps), and TFRs were averaged separately for time-windows marked as slow or fast by the algorithm. To visualize the spatial layout of spindles in the scalp EEG, density values per scalp electrode were averaged across participants, separately for the algorithmically detected slow and fast spindles.

### Thalamic and thalamocortical spindle co-occurrence

To evaluate whether spindles occurred in isolation or co-occurred across channels, we created continuous spindle time-courses across all channels. That is, a channel x time point matrix spanning the total duration of artifact-free NREM sleep was marked with absence or presence of spindles (0 or 1, respectively). The continuous spindle time-courses of a participant’s channels were divided into separate spindle events as follows: First, each time point was summed across channels. The resulting vector contained information on how many channels showed spindle activity at a given time point. Time points with 0 values meant no spindle activity occurring in any of the channels, and served to define boundaries between individual spindles. Time points with values > 0 were categorised as a (co-)occurring spindle.

We refer to the different combinations of thalamic and thalamocortical channels participating in a spindle event as spindle scenarios. Figure 2B schematizes these spindle scenarios at the thalamic (T spindles, 2B.I), the thalamocortical (TC spindles, 2B.II) and the cortical (T_0_C spindles, 2B.II) level. At the thalamic level, spindles were further categorized into clusters spanning 1 to 6 channels (T_1_ to T_6_ spindles, according to the number of thalamic channels involved in each spindle event). Spindle events occurring in >1 thalamic channel were further categorized as unilateral or bilateral, depending on whether they occurred in one hemisphere (unilateral; fig. 2B.I & II, second column) or in both (bilateral; fig. 2B.I & II third column). Because we recorded from 3 channels in each thalamic hemisphere, spindle clusters T_4_ or greater were bilateral by definition, while T_2_ and T_3_ could be either unilateral or bilateral clusters.

**Figure 2.**
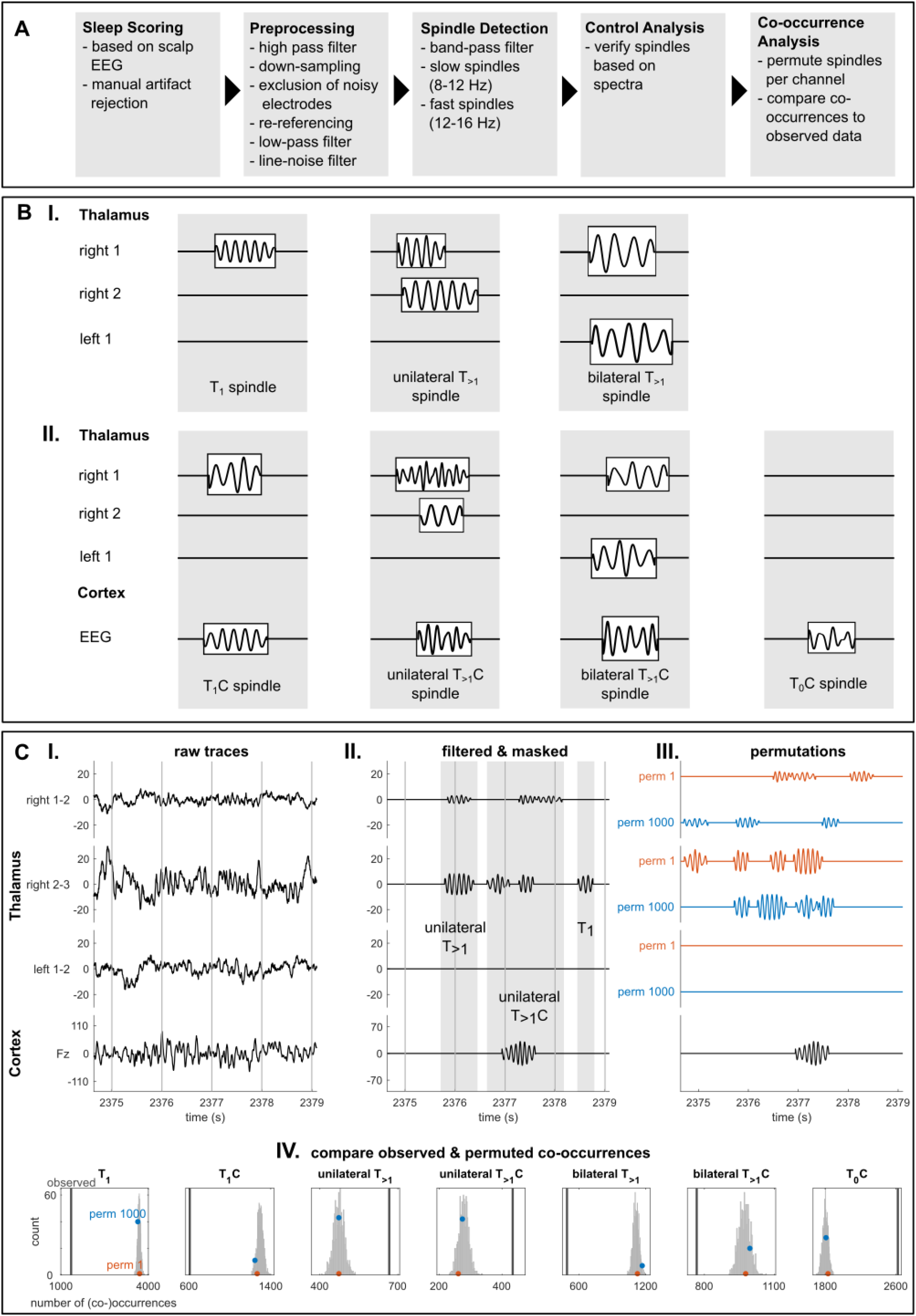
Overview of (pre-)processing steps to measure spindle (co-)occurrence in thalamic and scalp signals. **A)** Sequence of pre-processing steps and spindle detection. **B)** Schematized overview of spindle scenarios, i.e., possible channel combinations involved in a thalamic (**B.I**) or thalamocortical (**B.II**) spindle event. Three thalamic (2 right, 1 left) channels and the cortex time-course (all scalp EEG activity collapsed into one channel) are schematized. Subscripts describe the number of thalamic channels participating in a given spindle, so that a T_1_ spindle describes a spindle occurring in one thalamic channel. For more data examples for the spindle scenarios, see supplementary figure 1. **C)** Analysis steps are visualized on a data segment from three thalamic (2 right, 1 left) channels and one cortical channel (Fz) of one participant. Shown are an example raw trace (**C.I**) and filtered signal (**C.II**), in which only the detected spindles are shown and other data are masked. The grayscale columns mark (co-)occurring spindle activity and categorization of the shown spindles according to the spindle scenarios detailed in B). **C.III** depicts exemplar permutations of this data segment. Note that NREM time points were re-shuffled in each thalamic channel keeping the number and duration of spindles constant. The observed and permuted (co-)occurrences were then compared, as visualized in **C.IV**. Per spindle scenario, the number of co-occurrences were counted in the observed (black line) and permuted data (permutation 1 and 1000 are indicated in the histogram with the y-values at the lowest and highest bin-counts, respectively). To evaluate the likelihood of the observed value stemming from the permuted distribution, we assessed the proportion of permuted values that were more extreme than the observed value. In the analysis of thalamic and thalamocortical spindle clusters, we report the difference between the observed value and the mean of the permuted distribution. Note that the observed value and permuted distributions are shown as untransformed data for visualization purposes, but that we report percentages to enable comparison across subjects.

Given the difference in spindle numbers per scenario and participant, rates of each scenario are reported as the percentages rather than as raw counts. The analysis was carried out separately for thalamic and thalamocortical spindles. For thalamic spindles, we did not consider scalp EEG activity, and report the spindle events per thalamic channel as a percentage of all thalamic spindle events. For thalamocortical spindles, the information about spindle occurrence of all scalp EEG channels were collapsed into one cortical channel (Fig. 2B.II). That is, whenever a spindle occurred in one scalp EEG electrode, this was marked in the common scalp channel. Thalamocortical spindle events are reported as a percentage of all spindle events occurring across thalamus and cortex. We refer to thalamocortical spindles as T_x_C, where X correspond to the number of thalamus channels involved. In addition, we recorded how many scalp EEG spindle events did not have a thalamic counterpart as a percentage of the total number of detected spindles across all channels in a participant (T_0_C; Fig. 2B.II, fourth column). All analyses were computed separately for slow and fast spindles.

### Thalamocortical spindle co-occurrence maps

To evaluate putative topographical differences in thalamocortical spindle co-occurrence for the thalamic spindle scenarios (T_1_C, unilateral or bilateral T_>1_C), we focused on the continuous spindle time-courses across individual scalp EEG channels. To measure the co-occurrence of thalamic spindles with cortical spindles, we considered each thalamus channel as seed and each cortical channel as target channels. Only time points were included during which the seed and at least one cortical channel showed spindle activity. The spindle activity overlap was then calculated as the proportion of time points during which the target channel showed spindle activity at the same time as the seed channel out of the total number of T_1_C, unilateral or bilateral T_>1_C spindle time points of a seed channel. That is, in order to have 100% overlap with T_1_C spindles, the target channel would need to show spindle activity at all the time points that the seed channel was engaged in a spindle with no other thalamic channel showing simultaneous spindle activity. Consequently, a 50% overlap with T_1_C spindles would signify that only half of the target’s spindle activity temporally coincided with the seed’s spindle activity, when this particular thalamic seed channel showed a spindle that was not shared by any other thalamic channels. This analysis was carried out for each of the six thalamic channels, and separately for slow and fast spindles.

### Paired time-lag analysis of thalamocortical overlap

Thalamocortical overlap maps were calculated at a 0 time-lag, and thereby might not capture time-delayed relationships between seed and target channels. To evaluate whether there were any consistent non-zero time-lag relationships between seed and target channels across participants, histograms for the time lag between the onset of seed and target spindles were computed, sorted into bins of 0.05 s between ±0.75 s and averaged across seed, target channels and participants. The resulting histograms contain the average distribution of lags between thalamus and cortex, for which we tested the difference from 0 by means of a one-sample t-test.

### Within-participant statistical analyses

Spindle co-occurrence across channels may happen by chance according to the number and duration of spindles in a given channel pair. To test whether and to what extent observations of thalamic or thalamocortical spindle co-occurrence were due to chance, we tested within-participant statistical significance by permuting the data 1000 times. Figure 2C.III and IV visualizes this procedure. The time points of artifact-free NREM sleep of each thalamic channel were shuffled, keeping number and duration of spindles constant. Spindle co-occurrences in the permuted data were quantified as in the observed data, by using 0-columns in the channel x time matrix as bounds between spindle events (as shown in 2C.II). When permuting the data, T_>1_ spindles may be broken up into many T_1_ spindles, thus permuted data could contain a higher total number of spindle events than the original data. In order to keep permuted and non-permuted data comparable, we assessed the counts of spindle clusters as percentages of the sum of spindle events in a given participant and permutation. For the analysis of slow and of fast thalamic spindle clusters, we report the observed spindle events as the percentage of all respective spindle events across all thalamic channels. We repeated this analysis for slow and fast thalamocortical spindles separately, where we report the observed spindle events as the percentage of all respective spindle events across thalamic and cortical channels.

For statistical testing, the permuted and non-permuted data were z-scored based on the mean and standard deviation of the permuted data. The z-scored percentage of spindles in the non-permuted data were tested two-sided against the z-scored permuted distribution of percentage of spindles, with the p-value calculated as the proportion of permuted values that were more extreme than the observed value. For a visualization of the comparison between observed and permuted data, consider figure 2C.IV, which depicts exemplar permuted distributions and observed values for each of the thalamic and thalamocortical spindle scenarios for one participant. Lastly, the difference percentage of observed and mean permuted spindle clusters was computed. Positive values thus indicate cases in which specific spindle scenarios occurred more frequently in the observed than permuted data (e.g. 2C.IV, unilateral T_>1_C spindles), whereas negative values reflect spindle scenarios that occurred less frequently in the observed than permuted data (e.g., 2C.IV, T_1_ spindles). Consequently, scenarios that are more likely in the observed than permuted data suggest systematic temporal co-occurrence of spindle activity.

For mapping thalamocortical spindles, temporal spindle overlap was quantified based on the continuous spindle time-courses. Hence, the permuted data were not separated into spindle events, but quantified as the overlap of spindle time points of the target electrode with the seed electrode. The overlap thus describes the percentage of time a channel pair engages in spindle activity simultaneously. Only channels with p-value < 0.025 (indicating an observed value more extreme on either side of the permuted distribution) were considered in further analyses.

### Group level statistical analyses

Between-participant effects were tested by running repeated measures analysis of variance (RMANOVA) on the difference in spindle percentage between observed and permuted values. Where sphericity was violated, Greenhouse-Geisser corrected degrees of freedom are reported. For significant main and interaction effects, post-hoc pairwise comparisons were computed and adjusted for multiple comparisons using Bonferroni correction, and confidence intervals (Cis) at 95% are reported. The significance level was set at α = 0.05.

For analysis of thalamic spindle clusters, a RMANOVA was computed with factors *cluster size* (1…6 channels involved) and *spindle type* (slow vs. fast). For testing thalamic uni- vs. bilateral involvement, the RMANOVA factors were *cluster size* (2 vs. 3 channels involved), *hemispheric involvement* (unilateral vs. bilateral) and *spindle type* (slow vs. fast). To assess the thalamocortical spindle behavior, the following RMANOVA factors were defined: *thalamic involvement* (T_1_, unilateral or bilateral T_>1_), *cortical involvement* (T vs. TC spindles) and *spindle type* (slow vs. fast). EEG spindle events that did not have a thalamic counterpart were compared between slow and fast spindles, using paired samples t-tests.

To investigate thalamocortical maps across participants, we computed the median difference of observed from permuted thalamocortical overlap across thalamic seed channels, hemispheres and participants, but separately per *spindle type* (slow vs. fast spindles), and *thalamic involvement* (T_1_C, unilateral or bilateral T_>1_C). We only included datapoints that were significantly different from permuted data at the within-participant level.

To qualitatively compare the topographies of different spindle types and thalamic classes, we computed the center of gravity (CoG) per category. CoG quantifies the mean location of thalamocortical overlap, as the average x- and y-position of all electrodes weighted by their values of thalamocortical overlap. The x- and y-position are both relative to the central EEG electrode (Cz) and indicate lateralization and placement on the anterior/posterior axis, respectively. Given that thalamocortical maps were averaged across hemispheres, positive, zero and negative lateralization values describe ipsilateral, midline and contralateral locations, respectively. Positive, zero and negative y-values describe anterior, central and posterior locations, respectively. To test the robustness of estimating the CoG based on the average maps, we used a bootstrapping approach, in which we resampled the population of statistically significantly overlapping scalp EEG channels (n = 42), with replacement 1000 times. For each iteration, a new CoG was computed per condition (spindle type and thalamic involvement). Additionally, the difference between slow and fast spindles for each thalamic class was calculated for x and y coordinates, respectively.

Statistical testing was carried out to evaluate whether the CoG was significantly lateralized and/or shifted along the anterior/posterior axis for each of the thalamic classes and spindle types. For this, we tested the difference of each of the 6 bootstrapped distributions from 0, both for the x- and y-coordinates, by calculating the proportion of cases < 0 in each distribution. Next, we evaluated whether the difference of lateralization and/or shift along the anterior/posterior axis differed between slow and fast spindles, by calculating the proportion of cases < 0 in the distribution of the difference.

### Data availability

The data sets analyzed in the current study are not publicly available due to privacy concerns as outlined in the consent form.

## Results

To investigate thalamocortical spindle co-occurrence we analyzed data recorded from 7 participants during one night of sleep. On average, participants slept for 8.15 ± 1.55 hours (489 ± 93 minutes), of which 5.98 ± 1.46 hours (359 ± 87.33 min) were spent in NREM sleep (stages N2 and N3; see table 2 for participant-specific details of minutes spent in each sleep stage). We algorithmically detected slow and fast spindles in a total of 42 bipolar thalamic channels (6 per participant) and 133 scalp EEG electrodes (19 per participant). Note that because electrode placement into the specific subnuclei varied inter-individually, we assessed spindles in the anterior thalamus as a whole rather than its subnuclei. Accordingly, we generally refer to the anterior thalamus as the broad region. In contrast, we refer to the specific subnucleus as the anterior nucleus of the thalamus or ANT.

Table 2 reports the number of putative spindle events detected in the thalamic LFP and scalp EEG per participant, and the number of spindle events after the spindle control analysis. Only spindle events that passed the control analysis were included in all further analyses.

**Table 1.**
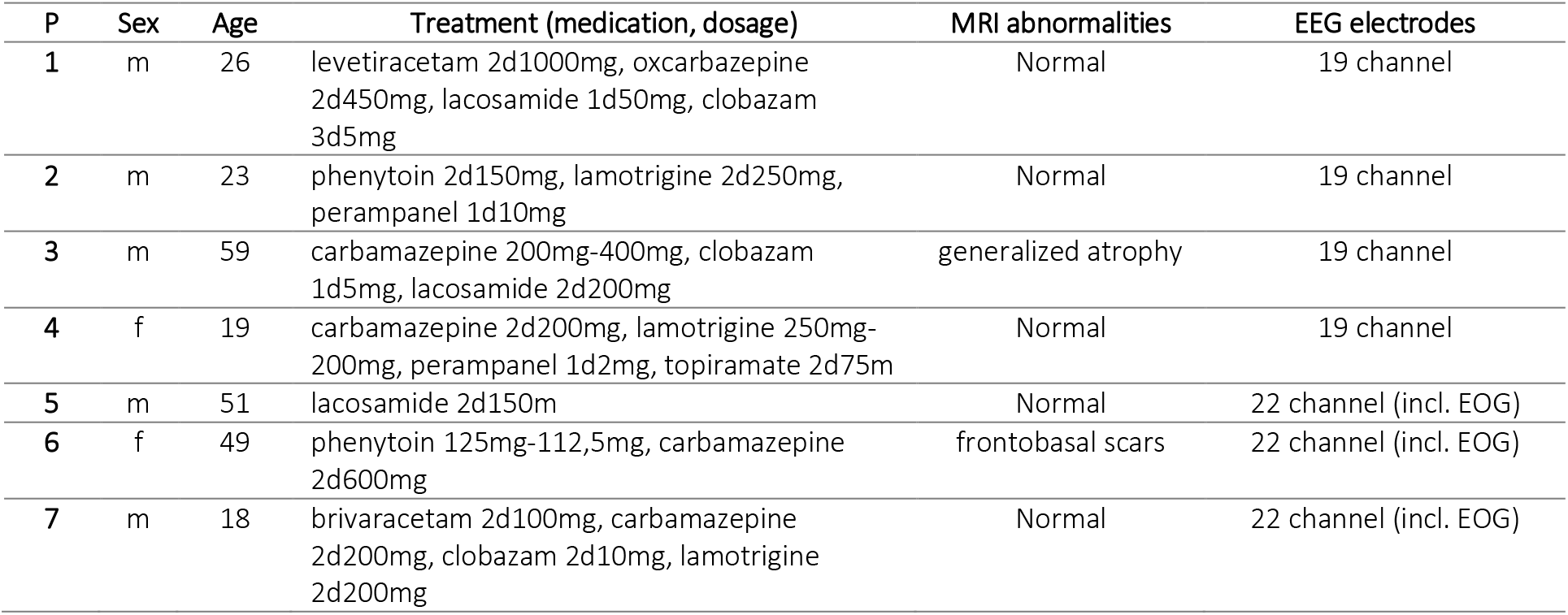
Individual clinical and MRI data and EEG setup.

| P | Sex | Age | Treatment (medication, dosage) | MRI abnormalities | EEG electrodes |
| --- | --- | --- | --- | --- | --- |
| 1 | m | 26 | levetiracetam 2d1000mg, oxcarbazepine 2d450mg, lacosamide 1d50mg, clobazam 3d5mg | Normal | 19 channel |
| 2 | m | 23 | phenytoin 2d150mg, lamotrigine 2d250mg, perampanel 1d10mg | Normal | 19 channel |
| 3 | m | 59 | carbamazepine 200mg-400mg, clobazam 1d5mg, lacosamide 2d200mg | generalized atrophy | 19 channel |
| 4 | f | 19 | carbamazepine 2d200mg, lamotrigine 250mg-200mg, perampanel 1d2mg, topiramate 2d75m | Normal | 19 channel |
| 5 | m | 51 | lacosamide 2d150m | Normal | 22 channel (incl. EOG) |
| 6 | f | 49 | phenytoin 125mg-112,5mg, carbamazepine 2d600mg | frontobasal scars | 22 channel (incl. EOG) |
| 7 | m | 18 | brivaracetam 2d100mg, carbamazepine 2d200mg, clobazam 2d10mg, lamotrigine 2d200mg | Normal | 22 channel (incl. EOG) |

**Table 2.**
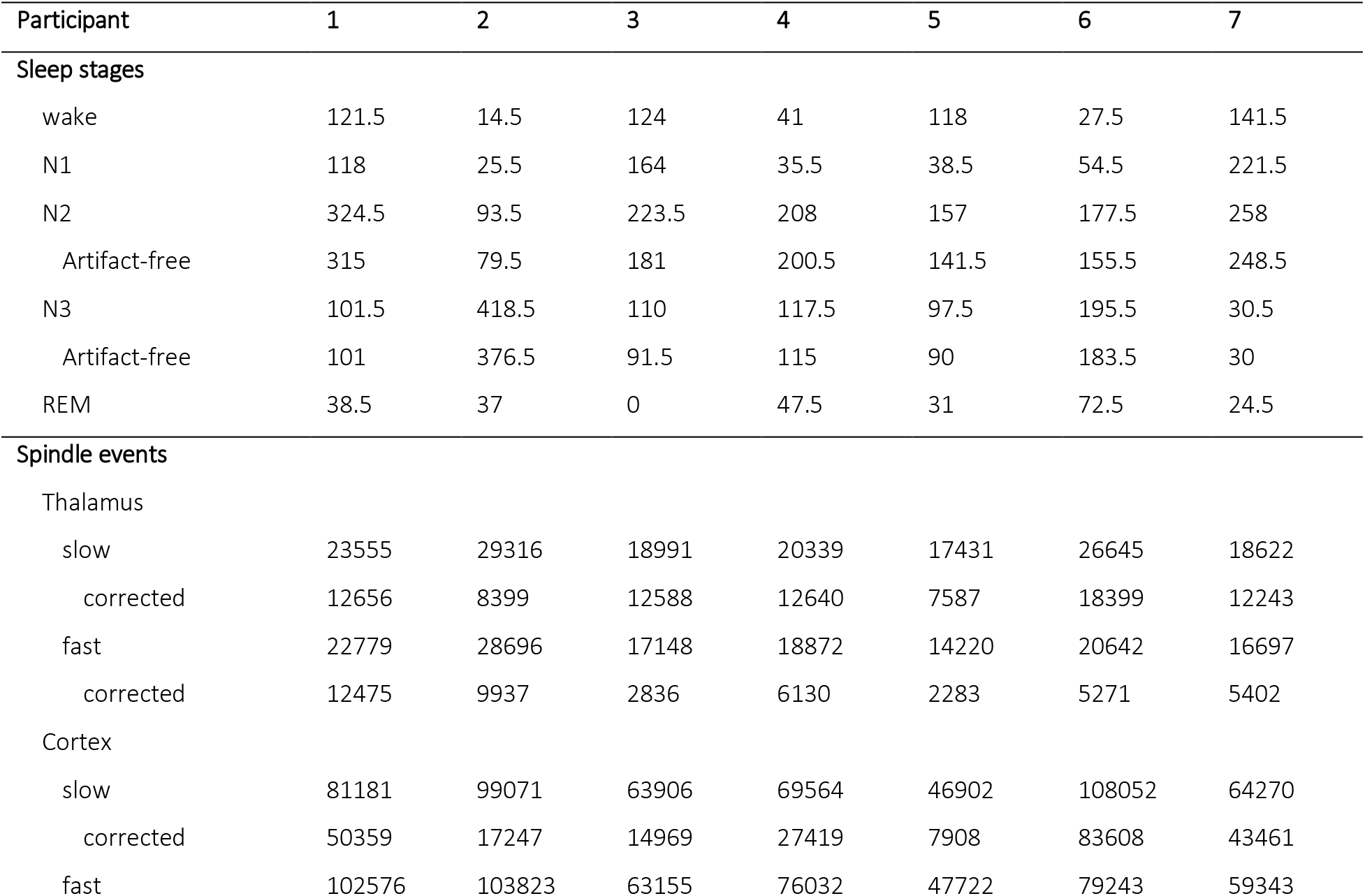
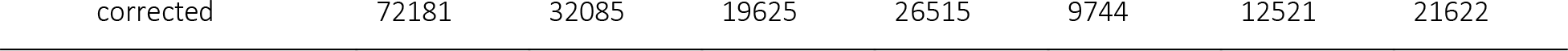
Overview of participants’ sleep architecture and detected spindle events before and after control analysis (corrected). Sleep staging data are reported in minutes, for spindle events the total number of events are reported (pooled across thalamic or scalp EEG electrodes, respectively).

### Slow spindles occur more frequently in anterior thalamus than fast spindles

We first investigated whether slow and fast spindles were present in the anterior thalamus and scalp EEG. Both slow and fast spindles were present in all participants. Figure 3 A shows an example of a slow followed by a fast spindle in one participant (for another example, see supplementary figure 1 B). The average TFRs of all slow and fast spindles in the same channel are shown in figure 3B. Note that the TFRs were computed across the slow and fast spindle band. Consequently, if the distinction into slow and fast spindles were arbitrary, this should be seen by an overlap in the TFR. Figure 3C depicts the average slow and fast spindle distributions in the scalp EEG across all participants.

**Figure 3.**
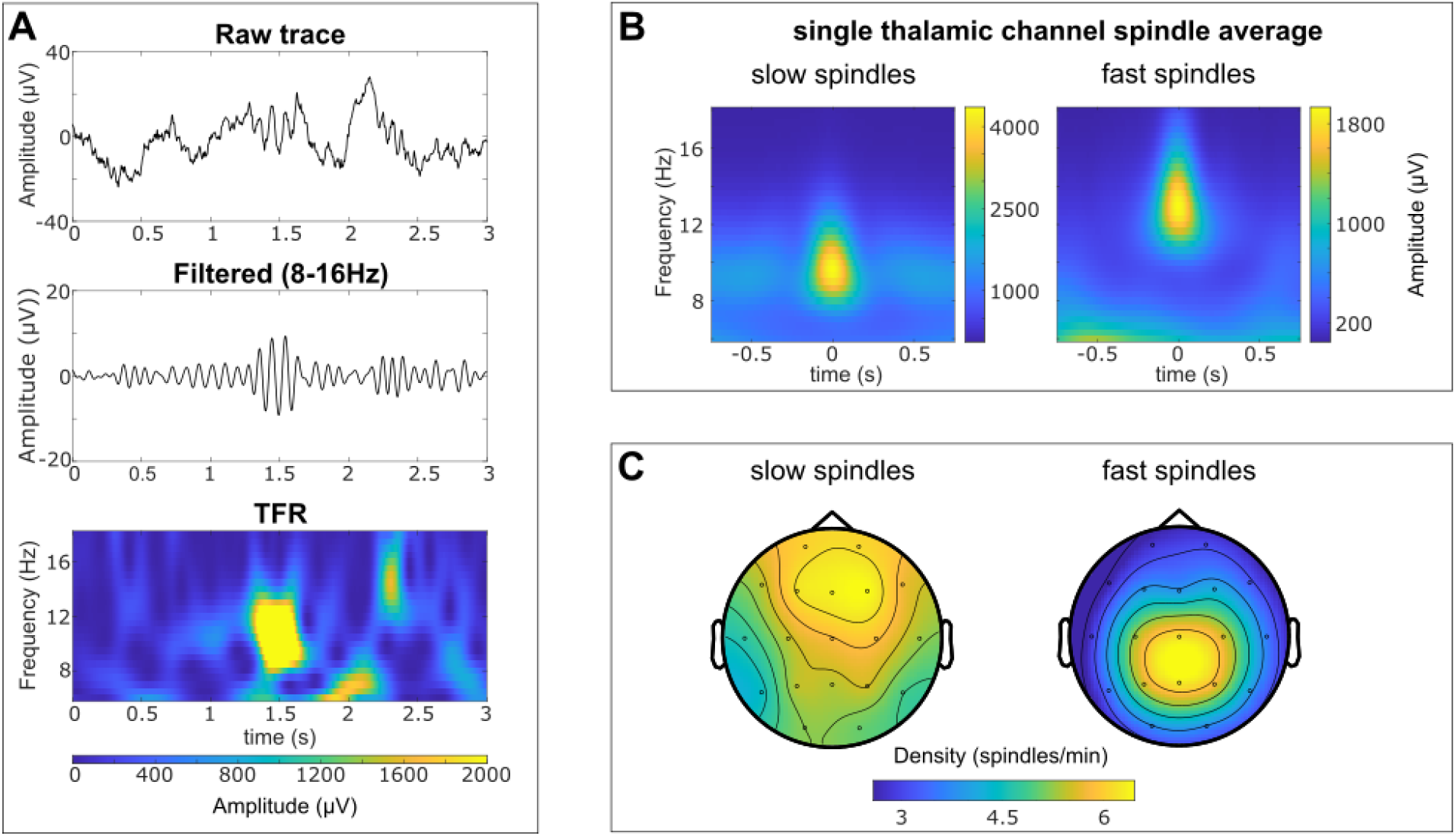
Slow and fast spindles in anterior thalamus and scalp EEG. **A)** Data of a slow (at 1.5 s) and fast (at 2.25 s) thalamic spindle, here represented in the raw trace (top panel), filtered trace (middle panel) and time-frequency representation (TFR; bottom panel). Data are shown for right thalamic channel 2-3 of participant 3. **B)** Averaged TFRs centered on time windows containing slow (left) and fast (right) spindles in one thalamic channel. The represented channel is the same as in A. **C)** Density of slow (left) and fast (right) spindles in scalp EEG averaged across all subjects.

Next, we assessed whether anterior thalamic spindles differed in their density, duration or amplitude. Figure 4 shows the spindle density (A), duration (B) and amplitude (C) for slow and fast spindles. Both data pooled across thalamic channel per participant (top row) are shown, as well as the histograms of the differences (slow – fast spindles) at the channel level (bottom row). On average, spindle density (fig. 4A) was higher (t(41) = 5.52; p < 0.001) for slow (6.34 ± 3.22 spindles/min) than fast spindles (3.01 ± 1.46 spindles/min). Spindle duration (fig. 4B; t(41) = 1.45; p = 0.15) and spindle amplitude (fig. 4C; t(41) = 1.97; p = 0.06) did not differ between slow and fast spindles.

**Figure 4.**
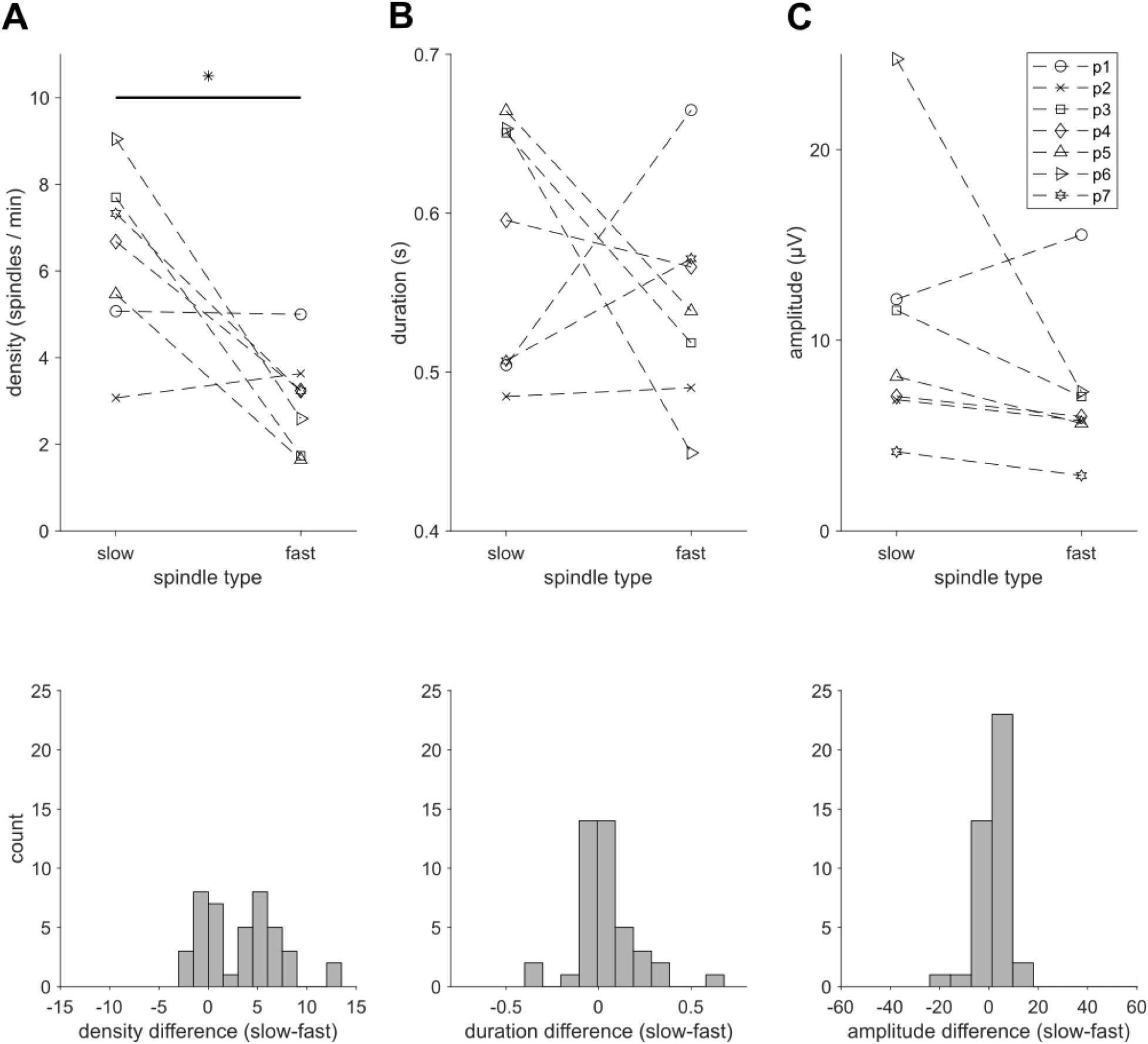
Spindle characteristics in anterior thalamus. Electrodes pooled per participant are shown (p1 to 7, legend) for slow and fast spindles (top row) and histograms of the differences (slow-fast) per electrode (bottom row). **A)** Average spindle density (number of spindles per minute) was on average higher for slow than for fast spindles; **B)** Spindle duration and **C)** spindle amplitude (peak to trough difference) did not differ between slow and fast spindles. Asterisk marks paired t-test difference significant at p < 0.05 (uncorrected).

We evaluated the same descriptive spindle measures when categorizing the data according to their corresponding estimated thalamic nuclei (suppl. material and suppl. fig. 1) and did not find any differences between thalamic nuclei.

### Characterization of spindles in the anterior thalamus

In a next step, we investigated whether slow and fast spindles co-occurred across channels in the thalamus (without considering the cortical EEG channels). We refer to the number of thalamic channels involved in a spindle as a channel cluster. Figure 5A depicts the channel cluster size of spindle events as a proportion of all thalamic spindles, separately for slow (upper panel) and fast (lower panel) spindles. To control for random spindle co-occurrence, we next ran permutations of the spindle time courses, with number and duration of spindles at each channel kept constant. These permuted data allowed us to estimate how often spindles occurred in isolation or co-occurred by chance. We compared these observed distributions of spindle clusters with the permuted distributions. Figure 5B shows the population difference between observed and permuted data for all 6 channel clusters for slow and fast spindles, respectively. Since the spindle cluster sizes are expressed in percentages, the likelihoods of T_1_ and T_>1_ clusters are interdependent and should be interpreted accordingly. T_1_ spindles, which were the most dominant type in the observed data (cf. fig. 5A), were less likely to occur in the observed compared to the permuted data. In contrast, spindle co-occurrences (T_>1_) occurred more often than expected by chance (F_1.29, 7.71_ = 119.41; p < 0.001 in RMANOVA; mean differences of observed – chance values between spindle cluster of T_1_ spindles vs. T_2…6_ channels range from −26.65 to −35.13 %; p ≤ 0.001). This suggests that thalamic spindles co-occur more often than would be assumed based on permuted distributions. Slow and fast spindles did not co-occur differently in the anterior thalamus (F_1,6_ = 0.18; p = 0.69).

**Figure 5.**
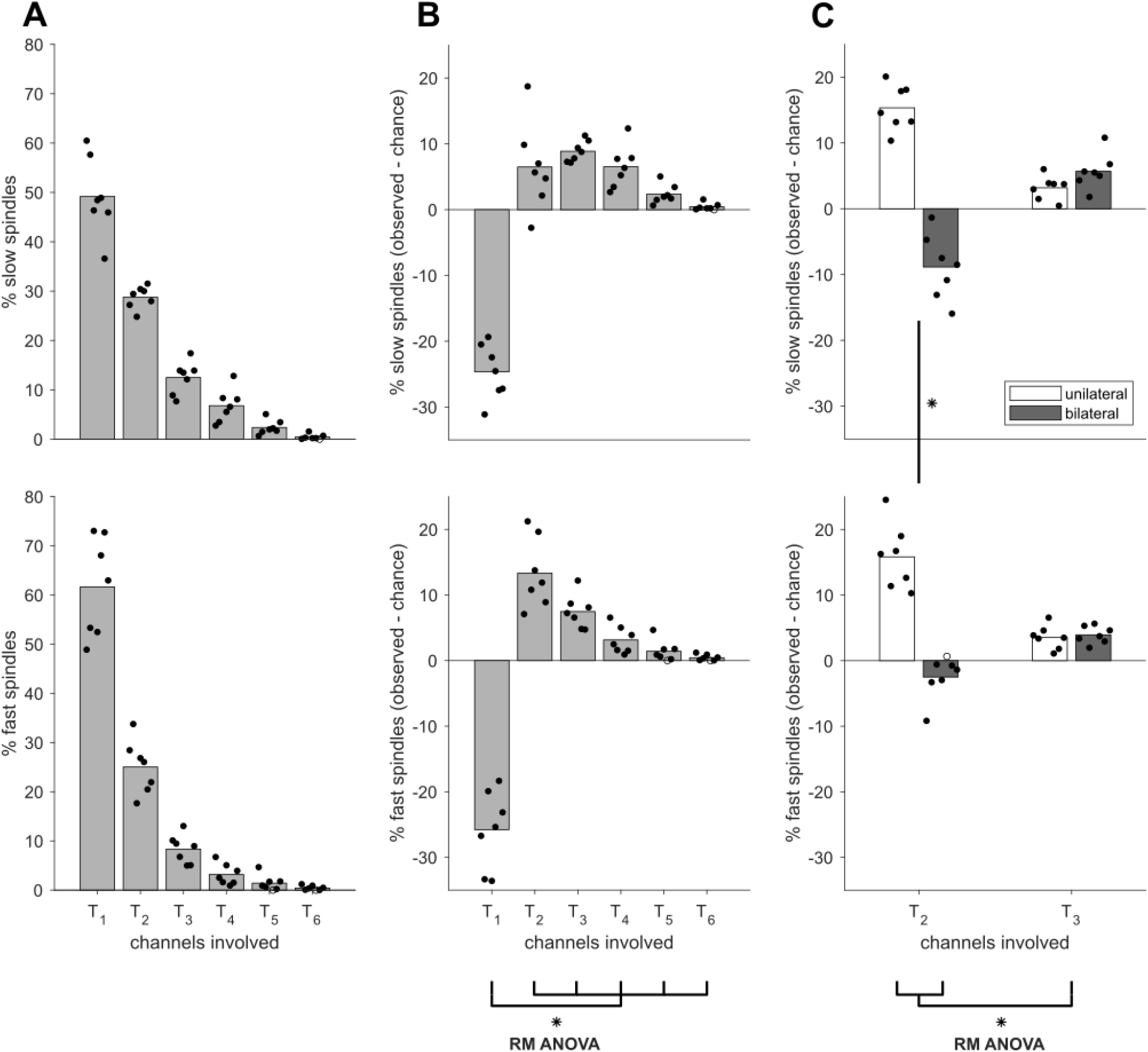
Observed thalamic spindles and the number of thalamic channels they spanned (channel clusters T_1_…T_6_) for slow (top) and fast (bottom) spindles. **A)** Observed percentages for each channel cluster out of the total number of slow or fast thalamic spindles, respectively. **B)** Difference of observed percentages from permuted data for thalamic spindle co-occurrences. Positive values demarcate when observed data are more likely to occur than by chance, negative values less likely than by chance. **C)** T_2_ and T_3_ spindles further subdivided into unilateral and bilateral slow and fast thalamic spindles. The difference of observed and permuted data is shown. Bars depict the group average, dots the individual participant data. Individual datapoints are systematically offset so that horizontal order corresponds to participants 1 to 7. Filled in dots are significantly different from permuted data at the within-participant level at p < 0.05 (two-sided), open dots depict data points failing to meet significance at the within-participant level. The asterisks mark differences significant at p < 0.05 in the RMANOVA (corrected for multiple comparisons), brackets stand for differences between means, forking out brackets for differences between individual bars.

It is noteworthy that some spindle events spanned more than three channels, pointing to the bilateral involvement of the thalamus in these spindle events (since our setup included three channels in each thalamic hemisphere). We subdivided the observed T_2_ and T_3_ clusters to investigate whether spindle clusters of two or three channels involved one or both hemispheres of the thalamus. Figure 5C depicts the difference in proportion of unilateral and bilateral T_2_ and T_3_ spindles between observed and permuted data, for slow and fast spindles, respectively.

T2 and T3 spindles showed distinct unilateral and bilateral involvement (F_1,6_ = 128.17; p < 0.001 in RMANOVA). We observed that unilateral T_2_ spindles occurred more often than by chance (15.58 %; 95 % CI, 11.99 to 19.18), while bilateral T_2_ spindles occurred less often than by chance (−5.68 %; 95 % CI, −7.46 to −3.89). This can also be seen in figure 5C when looking at the T_2_ data. This difference between unilateral and bilateral spindles was absent for T_3_ spindles: in this situation, both unilateral (3.36 %; 95 % CI, 1.95 to 4.77) and bilateral spindles (4.81 %; 95 % CI, 3.32 to 6.31) occurred equally more often than by chance (cf. fig. 5C, both unilateral and bilateral T_3_ bars are positive). We also observed a difference in the amount of slow and fast spindles spanning 2 or 3 channels in the thalamus (F_1,6_ = 7.88; p = 0.03 in RMANOVA). There were more fast T_2_ spindles (6.66 %; 95% CI, 4.21 to 9.12) than slow T_2_ spindles (3.24 %; 95% CI, 0.14 to 6.34), but equally many slow (4.44 %; 95% CI, 3.70 to 5.18) and fast T_3_ spindles (3.74 %; 95% CI, 2.56 to 4.93).

To conclude, these analyses indicated that compared to the permuted data, T_2_ spindles tended to be unilateral, whereas T_3_ spindles were equally likely unilateral or bilateral. Overall, it seems that thalamic spindle events did not only co-occur across channels, but were temporally coordinated within and across thalamic hemispheres.

### Multi- (but not single-) channel thalamic spindles show above-chance cortical counterparts

Next, we investigated whether observed thalamic spindle activity was systematically related to cortical spindle activity. For this analysis, spindles were subdivided according to the extent of thalamic involvement into T_1_, unilateral T_>1_ or bilateral T_>1_ spindles. We then categorized whether these spindles occurred solely in the thalamus (T spindles) or also had a cortical counterpart (TC spindles). We compared the occurrences of each spindle scenario to the total number of recorded spindles, rather than the total number of thalamic spindles, which was the case in the analysis up until now. Figure 6 shows the observed spindle percentages (A), and the difference between observed and permuted data (B), both for slow and fast spindles in the top and bottom row, respectively.

**Figure 6.**
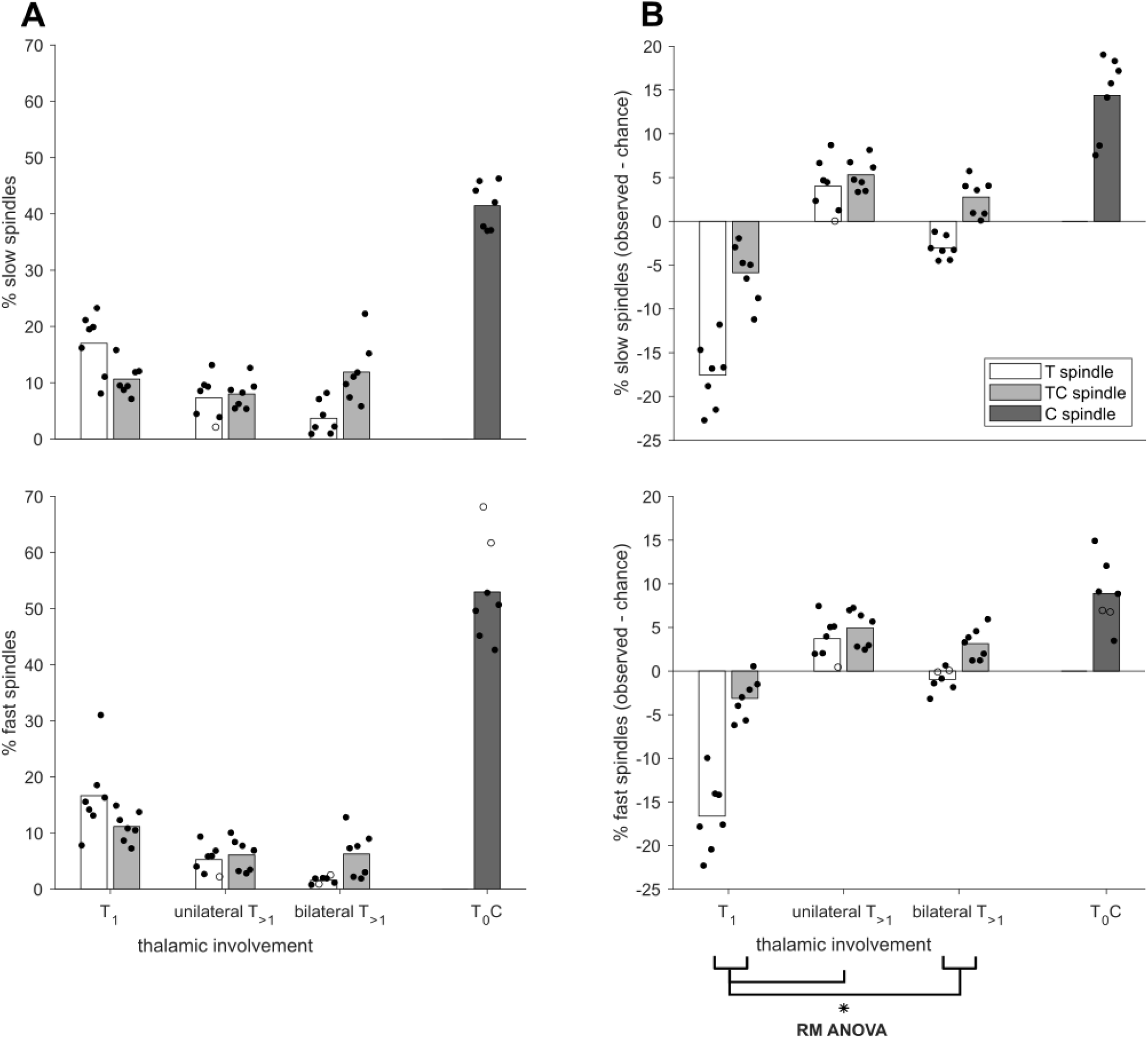
Observed spindles in thalamus and cortex, subdivided according to the level of involvement in the thalamus (T_1_ spindles or T_>1_ spindles, unilateral and bilateral), and whether they were accompanied by a cortical spindle (TC spindle, light grey) or not (T spindle, white). Cortical spindles without thalamic counterparts are also shown (T_0_C spindles, dark grey). **A)** Observation of each scenario as a percentage of the total number of slow (top) and fast (bottom) spindles. **B)** Difference of observed from permuted percentages for slow (top) and fast (bottom) spindles. Bars depict the group average, dots the individual participant data. Individual datapoints are systematically offset so that horizontal order corresponds to participants 1 to 7. Filled in dots are significantly different from permuted data at the within-participant level at p < 0.05 (two-sided), open dots depict data points failing to meet significance at the within-participant level. The asterisks mark differences significant at p < 0.05 in the RMANOVA (corrected for multiple comparisons), brackets stand for differences between means, forking out brackets for differences between individual bars. Note that there was no difference between slow and fast spindles.

T and TC spindles were more or less likely than in the permuted data, depending on whether they occurred in one channel or were unilateral or bilateral (F_1.04,6.24_ = 24.74; p = 0.002 in RMANOVA). While most observed slow and fast spindles involving the thalamus were T_1_ or T_1_C spindles (fig. 6A, most leftward bars), these types of spindles were less likely than expected by chance (cf. 6B, most leftward bars have negative values). Unilateral T_>1_ (3.87 %; 95 % CI, 1.70 to 6.05) and T_>1_C (5.12 %; 95 % CI, 3.73 to 6.51) spindles were more likely to occur than expected by chance. In contrast, bilateral T_>1_ spindles occurred less often than predicted (−2.0 %; 95 % CI, −2.80 to −1.20), whereas bilateral T_>1_C spindles occurred more often than predicted (2.96 %; 95 % CI, 1.30 to 4.67). These observations did not differ between slow and fast spindles (F_1.03,6.18_ = 0.62; p = 0.46 in RMANOVA).

Except for bilateral T_>1_ spindles without a cortical counterpart, spindles spanning more than one thalamic channel were more likely to occur than expected by chance. These multichannel spindles were also more likely to co-occur with cortical spindles. This can be seen in figure 6B, when looking at the unilateral and bilateral spindles (middle left and middle right bars). These results suggest that spindles at the thalamic (except for bilateral T_>1_ spindles) and thalamocortical level tend to co-occur across multiple channels systematically, indicating coordinated spindle activity.

We separately confirmed the absence of a systematic temporal offset between thalamic and cortical spindles (supplementary material & suppl. fig. 3). A temporal shift would bias analyses of spatiotemporal spindle overlap toward channel pairs with a smaller temporal offset, but this was not the case.

### Not all cortical spindles have an anterior thalamic counterpart

The data in the present study were recorded from a subsection of the thalamus and may have only captured a subsample of spindles occurring in the thalamus. Because cortical spindles are thought to depend on thalamic spindles and, more specifically, the projections of the different thalamic nuclei to distinct cortical networks, our data may also contain cortical spindles that did not have a thalamic counterpart (T_0_C spindles). To test this, we counted the occurrences of cortical-only spindles out of all recorded spindles, and compared them to permuted data. The most rightward bar in each panel of figure 6 shows the percentage of these T_0_C spindles. Indeed, in the observed data (fig. 6A), around half of observed cortical EEG spindles did not have a thalamic counterpart (slow: 41.46 ± 4.12 %; fast: 53.0 ± 9.04 %). This observation was higher than expected by chance (slow: 14.38 ± 4.60 %; fast: 8.88 ± 3.74 %), but did not differ between slow and fast spindles (t(6) = 1.87; p = 0.11). It may be that these T_0_C spindles had counterparts in thalamic subparts not sampled in our setup, such as more posterior nuclei.

### Patterns of thalamocortical spindle overlap depend on level of thalamic involvement and spindle subtypes

Lastly, we investigated cortical topographies of spindles dependent on the type of thalamic spindle and thalamic channel involvement. If coordinated thalamocortical spindle activity were to support targeted engagement of thalamocortical networks, different cortical spindle topographies should be observed based on different thalamic spindle properties. To compare differences in topography for thalamocortical spindles, depending on the spindle type (slow vs. fast) and the level of thalamic involvement (T_1_C, unilateral T_>1_C or bilateral T_>1_C spindles), we computed the thalamocortical overlap between all thalamic and cortical channel pairs across participants. The overlap quantifies the percentage of data points during which two channels (here, a seed channel in the thalamus and a target cortical channel) show simultaneous spindle activity. Since we verified that thalamocortical spindle onset occurred predominantly simultaneously (suppl. fig. 3), this analysis was carried out at 0-lag.

Figure 7 shows the median group difference of observed from permuted data for each category. The median group data were calculated based on the individual data that differed significantly from chance levels. The resulting topographies do not depict overlap in the left- and right hemisphere, but rather contra- or ipsilateral to the thalamic source, respectively, because data were averaged across participants, thalamus channels and hemispheres. Visual inspection suggests that thalamocortical overlap for slow spindles had a frontal topography (fig. 7 A-C), and fast spindles had a more central topography (fig. 7 D-F). Furthermore, the topographies seem to progress from lateralized to more widespread as more thalamic channels were involved in a thalamocortical spindle, which can be seen when comparing T_1_C (fig. 7 A & D) to unilateral (fig. 7 B&E) and bilateral (fig. 7 C&F) T_>1_C spindles, respectively.

**Figure 7.**
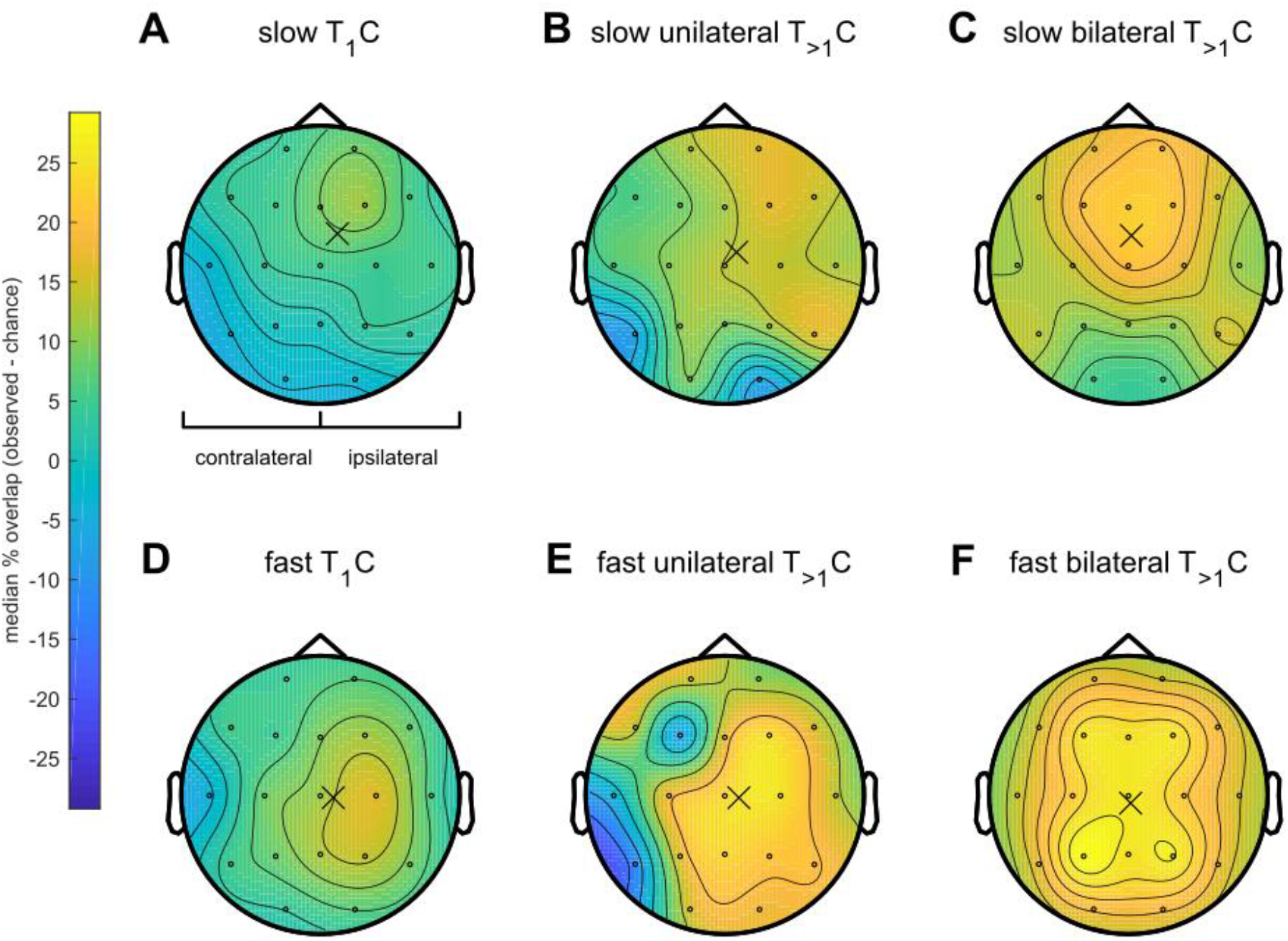
Thalamocortical spindle topographies. The difference of observed and permuted temporal overlap is shown as the group median. Accordingly, positive values describe a higher than by chance overlap, negative values a lower than by chance overlap. Data are averaged across thalamic channels, participants and hemispheres, such that values on the left within each topoplot describe the intensity of overlap contralateral and values on the right ipsilateral to the thalamus seed. The Center of Gravity is marked with a cross in each topoplot. Data are subdivided into spindle type and thalamic involvement: **A)** slow T_1_C spindles; **B)** slow unilateral T_>1_C spindles; **C)** slow bilateral T_>1_C spindles; **D)** fast T_1_C spindles; **E)** fast unilateral T_>1_C spindles; **F)** fast bilateral T_>1_C spindles.

To summarize these different topographies, we computed the Center of Gravity (CoG), which provides the spatial average weighted by the value of overlap at each cortical position. The CoG is marked with a cross in each panel of figure 7. Note, that the CoG does not necessarily exactly match the location suggested by the color scheme, given the different ways of calculating CoG and interpolating the colored surface.

For statistical analysis, we split the CoG into its x- and y-position allowing for quantifying the thalamocortical overlap’s lateralization and shift along the anterior/posterior (a/p) axis relative to a central scalp EEG topography (Cz). We used bootstrapping to statistically evaluate the differences in CoG between conditions. Figure 8 shows the bootstrapped CoG data for lateralization (fig. 8 A-D) and a/p-shift (fig. 8 E-H), split into the different conditions of thalamic involvement and spindle type (slow spindles in blue, fast spindles in orange).

**Figure 8.**
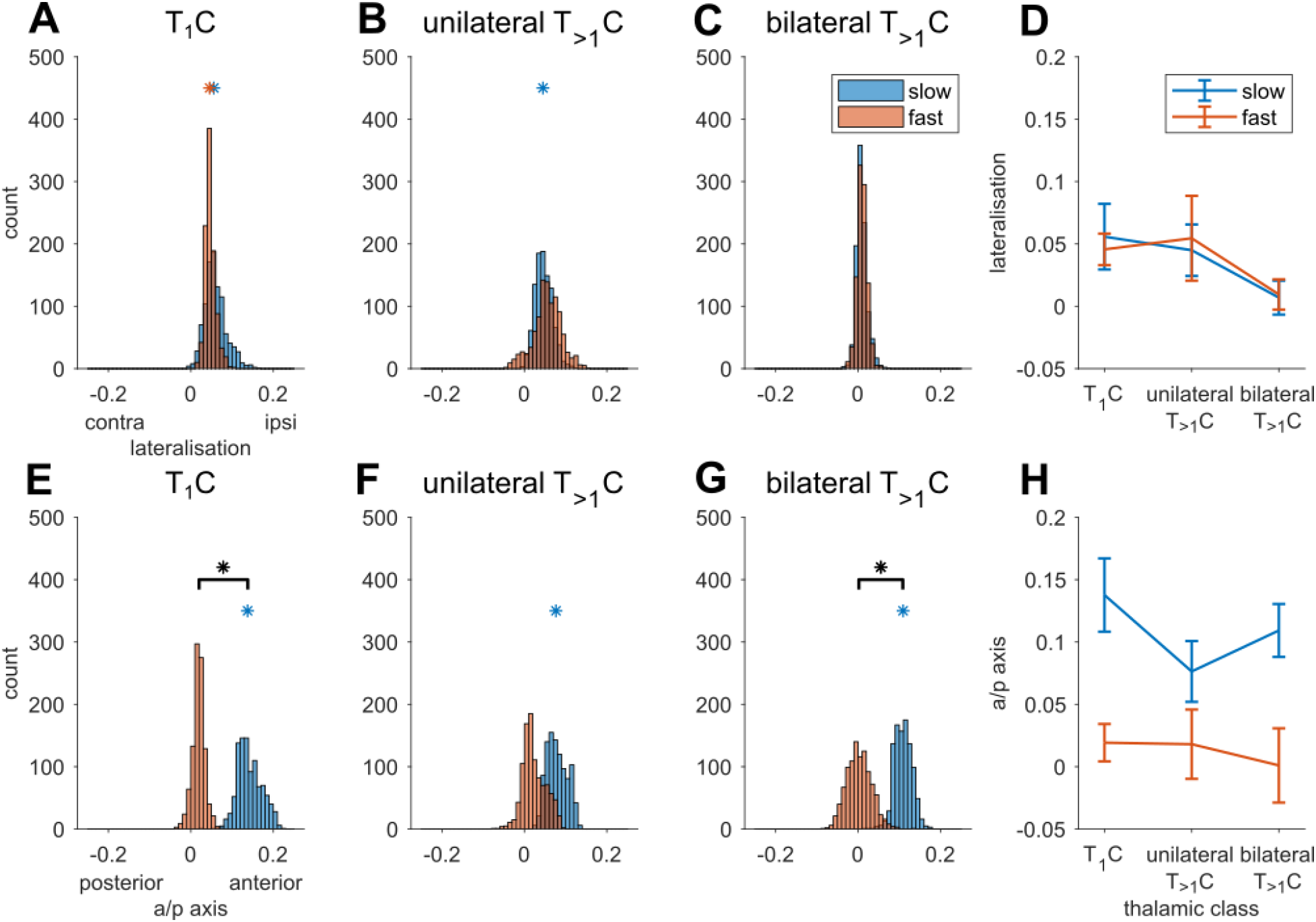
Center of Gravity of thalamocortical overlap, split into its x- and y-position, which describe lateralization and shift along the anterior/posterior (a/p) axis relative to electrode position Cz, respectively. Top row depicts the lateralization for **A)** T_1_C spindles; **B)** unilateral T_>1_C spindles; **C)** bilateral T_>1_C spindles; **D)** median values and standard deviations of the distributions in A-C. Bottom row depicts a/p shift for **E)** T_1_C spindles; **F)** unilateral T_>1_C spindles; **G)** bilateral T_>1_C spindles; **H)** median values and standard deviations of the distributions in E-G. Colored asterisks mark the distributions that differ from 0 at p < 0.05. Black asterisks and brackets mark distributions that differ from each other at p < 0.05.

Lateralization of spindles (fig. 8 A-C) was significantly different from 0 for slow T_1_C (0.055 ± 0.025; p = 0.001), slow unilateral T_>1_C (0.046 ± 0.02; p = 0.004), and fast T_1_C (0.045 ± 0.012; p = 0) spindles, indicating that the average thalamocortical overlap for these conditions tended to be focused ipsilateral to the thalamic spindles. Slow bilateral T_>1_C (0.008 ± 0.013; p = 0.224), as well as fast unilateral T_>1_C (0.055 ± 0.034; p = 0.095) and fast bilateral T_>1_C (0.01 ± 0.013; p = 0.19) spindles were not significantly lateralized. A/p-shift was significantly different from 0 for slow T_1_C (0.137 ± 0.029; p = 0), slow unilateral T_>1_C (0.075 ± 0.024; p = 0), and slow bilateral T_>1_C (0.108 ± 0.022; p = 0) spindles. Thus, the average CoG of thalamocortical spindle overlap of slow spindles was shifted anteriorly while fast T_1_C (0.018 ± 0.015; p = 0.102), fast unilateral T_>1_C (0.018 ± 0.028; p = 0.198) and fast bilateral T_>1_C spindles (0.004 ± 0.03; p = 0.471) were not shifted along the a/p axis.

When comparing slow and fast spindles, T_1_C slow spindles were significantly more anterior than T_1_C fast spindles (0.12 ± 0.033; p = 0) but equally lateralized (0.009 ± 0.028; p = 0.355). Slow and fast unilateral T_>1_C spindles did not differ in terms of lateralization (−0.009 ± 0.039; p = 0.585) or shift on the anterior/posterior axis (0.056 ± 0.037; p = 0.076). Slow bilateral T_>1_C spindles were significantly more anterior than fast bilateral T_>1_C spindles (0.106 ± 0.039; p = 0.008), but did not differ in their lateralization (−0.002 ± 0.018; p = 0.556).

These results point towards differences in thalamocortical overlap depending on the spindle type (slow vs. fast) and level of thalamic involvement. As such, we observed that slow spindles tended to co-occur with cortical slow spindles in frontal scalp EEG sites. Co-occurrence of thalamic and cortical fast spindles was distributed more widely across the cortex, which was reflected by a more centralized average of thalamocortical overlap. Additionally, the level of involvement of the thalamus in spindles was related to the cortical lateralization of thalamocortical overlap: During T_1_C spindles and slow unilateral T_>1_C spindles, cortical electrodes ipsilateral to the thalamus channel(s) tended to engage in spindles together, whereas bilateral T_>1_C spindles showed a higher overlap with bilateral cortical electrodes.

## Discussion

In the current study, we investigated the spatiotemporal coordination of spindles in the human anterior thalamus and cortex. Both slow and fast spindles were detected in the anterior thalamus, with a higher density of slow spindles. We categorized spindles based on their main frequency (slow vs. fast spindles) and level of thalamic involvement (spindles spanning one channel vs. multiple channels unilaterally vs. bilaterally) and compared observed patterns of spindle co-occurrence with permuted data. Figure 9 graphically summarizes predominant scenarios of thalamic and thalamocortical spindle (co-)occurrence we found in the permuted (A) and observed (B) data. We found that simultaneous multichannel spindle activity within thalamic and thalamocortical circuits occurred more often than expected by chance (fig. 9B). This points to spindle activity as a coordinated phenomenon across brain sites. Interestingly, multichannel spindles did not only co-occur unilaterally (fig. 9B.4 & 7), but also bilaterally within the thalamus (fig. 5; fig. 9B.5&6). When contrasting the different spindle subtypes based on their main frequency and the level of thalamic involvement, we observed similar rates of co-occurrence, but differences in the spatial pattern of cortical overlap (fig. 7). Overall, these results point to coordinated and differential patterns of sleep spindle co-occurrence in thalamocortical circuits.

**Figure 9.**
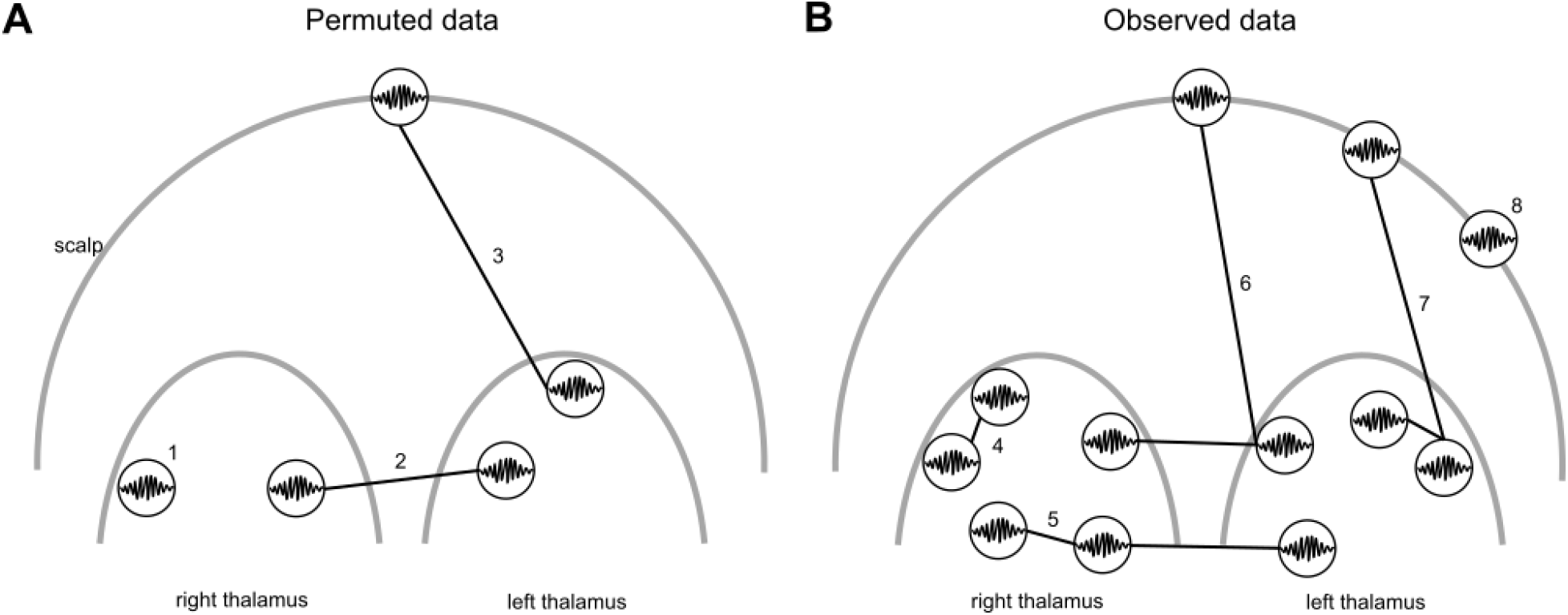
Graphical summary of thalamic and thalamocortical spindle (co-)occurrence, depicting spindle scenarios (numbered; cf. fig. 2 B) that appeared more often in the A) permuted or B) observed data. Bilateral thalamus and scalp are depicted schematically. Individual channels are indicated by circles, simultaneous spindle activity is indicated by connecting lines. **A)** In the permuted data, locally restricted spindles predominated. Thalamic spindles were predominantly T_1_ spindles (1) or bilateral T_2_ spindles (2), and thalamocortical spindles mostly T_1_C spindles (3). **B)** In the observed data, the most common spindle scenarios included two or more channels, indicating coordinated spindle activity. At the thalamic level, unilateral T_>1_ spindles (4) predominated, as well as bilateral T_>2_ spindles (5; here illustrated by a T_3_ spindle). Thalamocortical spindles were predominantly bilateral T_>1_C (6) or unilateral T_>1_C spindles (7). There were also more cortical spindles without a thalamic counterpart (T_0_C spindles; 8) than expected by chance.

There are alternative scenarios to coordinated and differentiated spindle activity across brain areas: For instance, spindles could be restricted to specific sites, which would preclude them from timing communication between brain areas during NREM sleep. Previous iEEG studies reported spindles as predominantly local in the cortex (Andrillon et al., 2011; Frauscher et al., 2015; Peter-Derex et al., 2012) and with low synchrony between cortical spindles and those in deeper brain structures like insula and hippocampus (Frauscher et al., 2015). These reports gave rise to the idea of sleep spindles as a local phenomenon. Although we observed many occurrences of spindles restricted to one channel, we also saw that spindles often spanned multiple channels, both at the thalamic (∼50 % of spindles) and the thalamocortical level (∼35% of all spindles engaged at least one thalamic and one cortical channel). Most of these multi-channel spindle scenarios occurred more often than predicted by chance, suggesting that spindle activity in thalamocortical networks is indeed coordinated rather than always being locally restricted. While we cannot exclude the possibility that some multichannel spindles in our data reflect volume conduction from a single source, volume conduction clearly cannot explain many of the patterns we observed, among which bilateral spindles picked up by electrodes in opposite hemispheres (cf. suppl.fig. 1C), or spindle onset and duration differences between neighbouring channels (cf. fig. 1C.I and II). Moreover, the proportion of multi-channel thalamic and thalamocortical spindles observed here, may be an underestimation as the exact thalamic recording sites differed between individuals, and consequently only sample a subsection of the thalamus. Events categorized here as T_1_, T_1_C or T_0_C spindles may have engaged thalamic regions not sampled in our setup. Our findings thus indicate that many spindles are not local, and do not co-occur merely by chance, but are coordinated over wide areas of the thalamocortical network.

Although pathological brains differ from healthy ones, treatment with benzodiazepines and anticonvulsant drugs may influence the sleep architecture (for a comprehensive review see Jain & Glauser, 2014) and spindles (Leong et al., 2022; Plante et al., 2015) and the wide age range in our sample may bias spindle-related characteristics (Muehlroth & Werkle-Bergner, 2020), clinical populations are presently the only opportunity to directly measure time-resolved recordings of the human thalamus. Previous studies have reported thalamocortical spindles in medial/anterior thalamus and cortex (Schreiner, Kaufmann, et al., 2021; Szalárdy et al., 2021) as well as posterior thalamus and cortex (Bastuji et al., 2020; Mak-McCully et al., 2017), albeit only one of these studies investigated thalamic and thalamocortical spindle co-occurrence in a more systematic fashion (Bastuji et al., 2020). In that study, around half of all thalamic spindles occurred in one channel and the remaining half spread across two or more channels. These reports were limited to the posterior thalamus due to clinical demands, were not contrasted with chance spindle co-occurrence and only investigated fast spindles. Our data now add information on the anterior thalamus.

In contrast to previous literature, which focused on fast spindles, we systematically investigated thalamic and thalamocortical spindle co-occurrence separately for slow and fast spindles. In our data, both spindle types were coordinated in a largely similar fashion, although some differences were also found. Slow and fast spindles were more likely to span multiple channels in the observed than permuted data, both unilaterally and bilaterally, and co-occurred between thalamus and neocortex. Our time lag analysis revealed simultaneous activation of thalamic and cortical spindles. The lack of temporal sequence in our data does not provide information on causal relations of spindle activity across regions. However, our findings provide evidence that both slow and fast spindles are present in the (anterior) thalamus, and that both engage in a coordinated fashion with the neocortex. It is thus not likely that slow spindles are specific to cortico-cortico interactions, as suggested previously (Mölle et al., 2011). Interestingly, slow and fast thalamocortical spindles did show different topographical patterns of thalamocortical overlap: slow thalamic spindles showed higher overlap with frontal scalp EEG electrodes, whereas fast thalamic spindles showed higher overlap with centro-posterior scalp EEG electrodes. These topographies of overlap are reminiscent of sites of slow and fast spindle activity observed in previous (i)EEG studies (Andrillon et al., 2011; Mölle et al., 2011; Peter-Derex et al., 2012). Note, however, that the different patterns we uncovered cannot be merely explained by the higher density of slow spindles in frontal EEG channels (cf. fig. 3C). We used permutation testing to control for the bias of higher spindle density/duration on spurious spindle overlap. Thus, although the frequency-based spindle dichotomy has been criticized as overly simplified (Gonzalez et al., 2021), our results suggest that thalamic spindles engage different cortical networks, depending on their main frequency.

The topographical patterns of thalamocortical spindle overlap were not only related to the main frequency of a given spindle, but also to the level of involvement of the thalamus. Andrillon and colleagues (2011) reported that spindles occasionally span multiple channels across the cortex. In this scenario, spindles travel along the cortical posterior-to-anterior axis (Andrillon et al., 2011; Muller et al., 2016). In the present study, we observed that multi-channel thalamic spindles were related to an increase in the number of cortical channels showing high thalamocortical spindle overlap. Consequently, the different cortical spindle scenarios (localized vs. global cortical events) reported in previous (i)EEG studies could be related to differential involvement of the thalamus in these spindles. Accordingly, multichannel cortical spindles could arise as a consequence of multichannel thalamic spindles. That is, when a spindle travels through the thalamus, this may be reflected as a cortical traveling spindle mediated by thalamocortical projections with the engaged thalamic sites. In fact, the patterns of thalamocortical spindle co-occurrence in our data (cf. fig. 7) suggest that this might be the case. For instance, the thalamocortical overlap of fast spindle events differed between spindles occurring in one thalamic channel and those spanning bilateral thalamic channels: In the former case, the thalamocortical overlap was spatially restricted to centro-posterior cortical sites ipsilateral to the thalamic site (cf. fig. 7 & 8), whereas in the latter case, the overlap was more widespread across the cortex bilaterally. Interestingly, cross-hemispheric co-occurrence of cortical spindles is preserved in callosotomized patients, suggesting that cortical spindle spread may depend on thalamocortical instead of cortico-cortico connections (Bernardi et al., 2021). The results of the present study add to the currently limited literature on interhemispheric spindle co-occurrence by showing that cortical spread of spindle activity is related to the level of involvement of the thalamus. More broadly, the involvement of different thalamic sites in a spindle may engage different cortical networks.

### Spindles may facilitate network communication during memory consolidation

Memory consolidation requires reactivation in hippocampo-cortical circuits to trigger structural changes (Josselyn et al., 2015). This reactivation must be specific to the neuronal populations involved in encoding the memory, and thus requires coordinated activity across (sub)cortical networks. Hippocampal ripples (80 – 140 Hz), which have been linked to memory reactivation (Zhang, Fell, & Axmacher, 2018), co-occur with spindles in the hippocampus and cortex (Ngo et al., 2020; Staresina et al., 2015), the anterior thalamus (Szalárdy et al., 2021) and in the anterior thalamus and hippocampus (Sarasso et al., 2014). Spindles are hypothesized to align windows of excitability in hippocampo-cortical circuits, and thereby coordinate the information transfer necessary for long term memory consolidation (Klinzing et al., 2019).

The thalamus is richly connected with cortical and subcortical structures (Cappe, Morel, Barone, & Rouiller, 2009; Jankowski et al., 2013; Zhang, Snyder, Shimony, Fox, & Raichle, 2010) and has been proposed as a connector hub between distinct cortical networks (Kawabata et al., 2021). Recent research hints at an active role of the thalamus in memory consolidation during sleep (Klinzing et al., 2019). According to this view, thalamus functions as a switchboard to enhance processing within and between specific cortical networks as required by task demands (Nakajima & Halassa, 2017; Schmitt et al., 2017). Those same mechanisms seem to be engaged during spindle generation in rodent NREM sleep (Chen et al., 2016; Halassa et al., 2014). Regarding memory reactivation, spindles in the thalamus (within or across thalamic nuclei) could selectively engage (sub)cortical sites through its rich projections. This could lead to carefully timed communication between the hippocampus and the cortical network(s) implicated in a specific memory and thereby enable reactivation of memory traces crucial for memory consolidation.

Previous studies have reported the importance of temporally (Latchoumane et al., 2017; Schreiner, Petzka, et al., 2021) and spatially (Petzka et al., 2022) organized cortical spindles during memory consolidation processes. Although our study did not include a memory task, here we add to this literature by showing that thalamocortical spindle activity is temporally coordinated and spatially specific. Further research using memory tasks needs to be conducted to directly study how the thalamocortical processes reported here may support memory consolidation.

## Conclusion

The present study investigated (co-)occurrence of sleep spindles in the anterior thalamus and neocortex. Up to half of the recorded spindles co-occurred across multiple channels, both at the thalamic and thalamocortical level. Although spindles were more likely to engage the thalamus unilaterally, we also observed simultaneous bilateral thalamic spindles. Slow and fast spindles were observed in the anterior thalamus. Both slow and fast, as well as single channel and multichannel thalamic spindles were associated with distinct topographical patterns of spindle co-occurrence in the cortex. As such, slow thalamocortical spindles generally co-occurred with more frontal scalp EEG channels, while fast thalamocortical spindles showed a wider cortical overlap, averaging highest at central scalp EEG sites. Single channel and unilateral thalamic spindles had more lateralized cortical co-occurrence patterns than bilateral thalamic spindles. These diverse patterns of thalamocortical spindle co-occurrence could be involved in selective engagement of thalamocortical networks during memory consolidation and other cognitive processes.

## Acknowledgements

The authors would like to thank all patients and their families for the cooperation in this study. In addition, the authors would like to thank Julio Rodriguez Larios for providing helpful feedback, and the editors and reviewers who have dedicated their time to provide comments and questions which greatly improved this manuscript.

## Author Contributions

**Hannah Bernhard**: Conceptualization, Data curation, Formal analysis, Methodology, Project administration, Resources, Software, Validation, Visualization, Writing – original draft, Writing – review & editing. **Fred Schaper**: Data curation, Investigation, Project administration, Resources, Visualization, Writing – review & editing. **Marcus Janssen**: Investigation, Resources, Writing – review & editing. **Erik Gommer**: Data curation, Investigation, Project administration, Resources, Writing – review & editing. **Bernadette Jansma**: Funding acquisition, Supervision, Writing – review & editing. **Vivianne Van Kranen-Mastenbroek**: Funding acquisition, Investigation, Resources, Supervision, Writing – review & editing. **Rob Rouhl**: Funding Acquisition, Investigation, Project administration, Resources, Supervision, Writing – review & editing. **Peter de Weerd**: Conceptualization, Funding acquisition, Methodology, Resources, Supervision, Writing – original draft, Writing – review & editing. **Joel Reithler**: Conceptualization, Formal analysis, Methodology, Resources, Supervision, Validation, Visualization, Writing – original draft, Writing – review & editing. **Mark Roberts**: Conceptualization, Formal analysis, Funding acquisition, Methodology, Resources, Software, Supervision, Validation, Visualization, Writing – original draft, Writing – review & editing. **Louis Wagner**: Investigation, Resources. **Albert Colon**: Investigation, Resources. **Danny Hilkman**: Investigation, Resources. **Marielle Vlooswijk**: Investigation, Resources. **Jeske Nelissen**: Investigation, Resources. **Linda Ackermans**: Investigation, Resources. **Yasin Temel**: Investigation, Resources.

## Supplementary material

**Supplementary figure 1.**
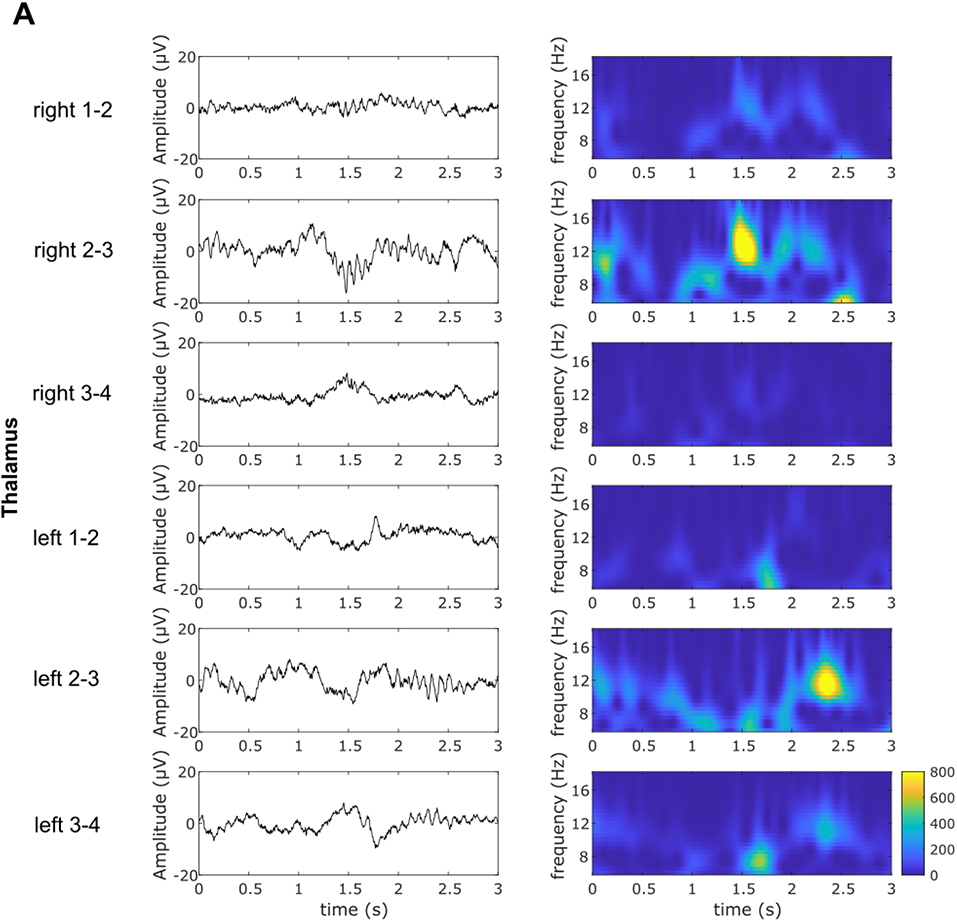
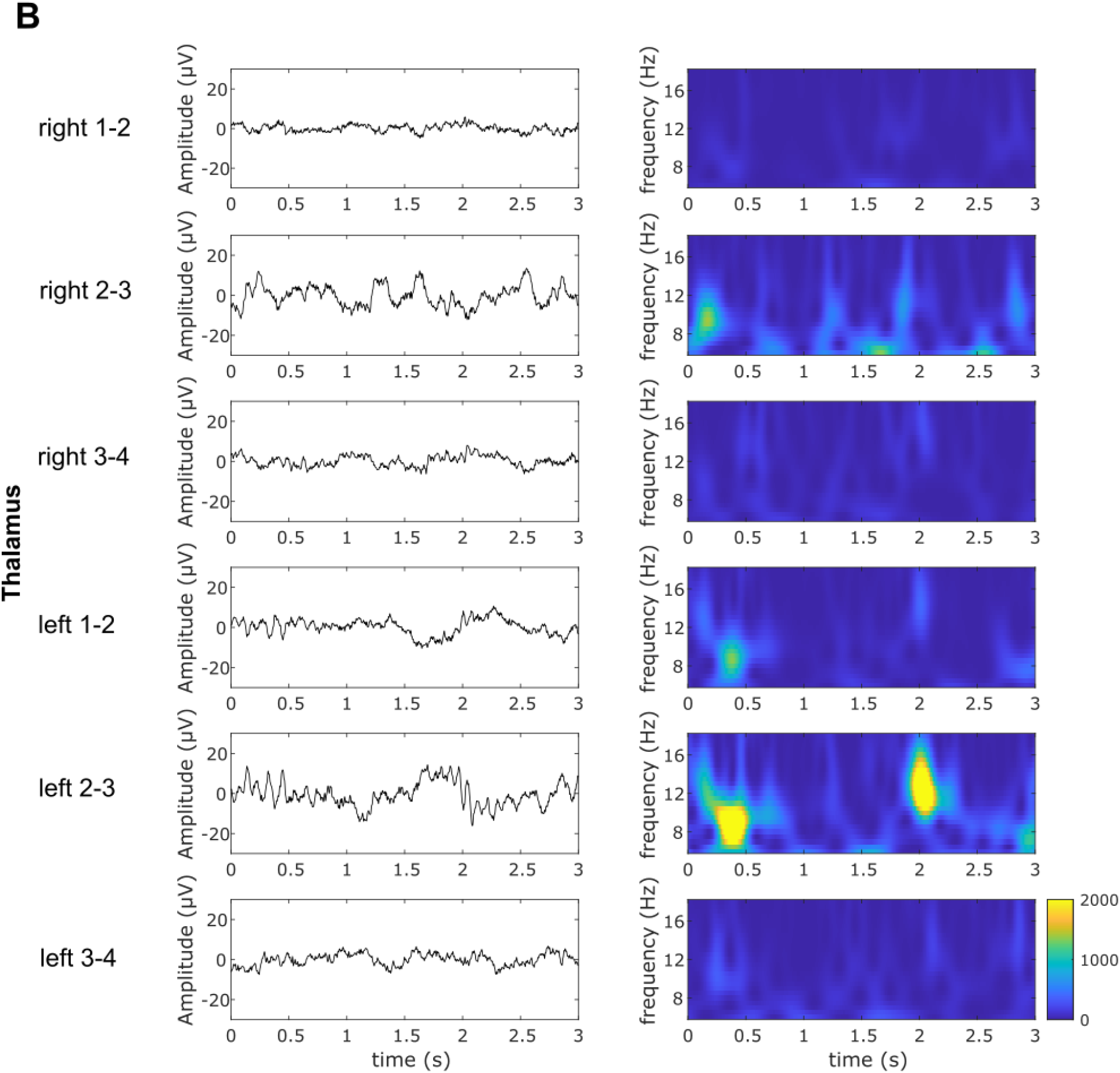
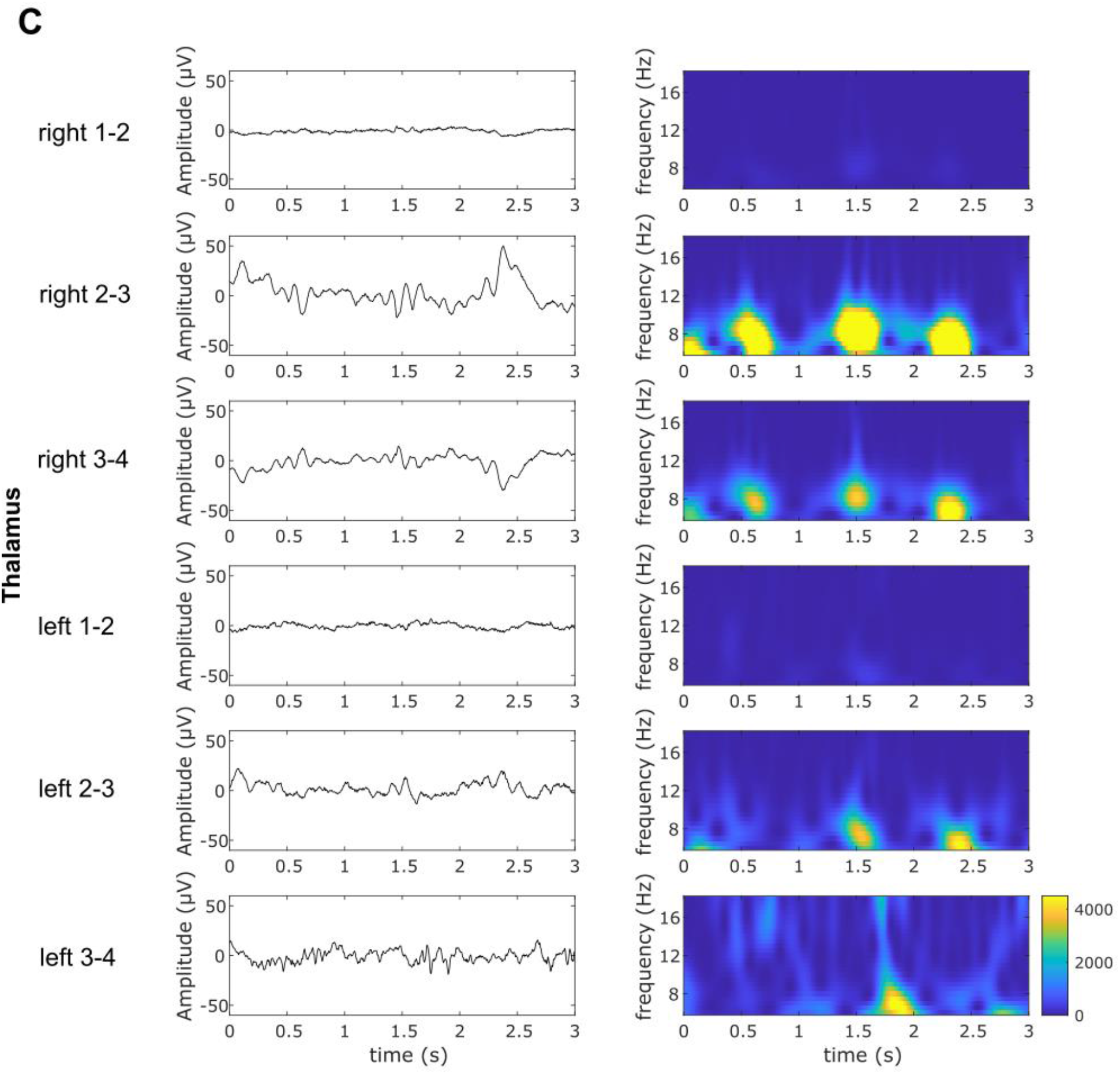
Representative data of thalamic spindles, shown in raw trace (left column) and time-frequency representations (TFRs; right column) from three different participants. **A)** A fast spindle in right thalamus (channel 2-3) at 1.5s is accompanied by a slow spindle in left thalamus (channel 3-4). This spindle is, in consequence, bilateral. At 2.3s, a unilateral spindle in the left thalamus occurs (channels 2-3 and 3-4). **B)** A slow bilateral spindle (channels right 2-3, left 1-2 and left 2-3) occurs at the beginning of the epoch. Though the peak of these spindles occurs at different time points, the tails of the spindles overlap, qualifying them as co-occurring spindles in our analysis. At 2s, we see a local fast spindle in channel left 2-3. **C)** A unilateral spindle in the right thalamus (channels 2-3, 3-4) at 0.5s, followed by bilateral spindles at 1.5s and 2.3s in the right (channels 2-3 and 3-4) and left (left 2-3, and partially left 3-4) thalamus.

**Supplementary Table 1.**
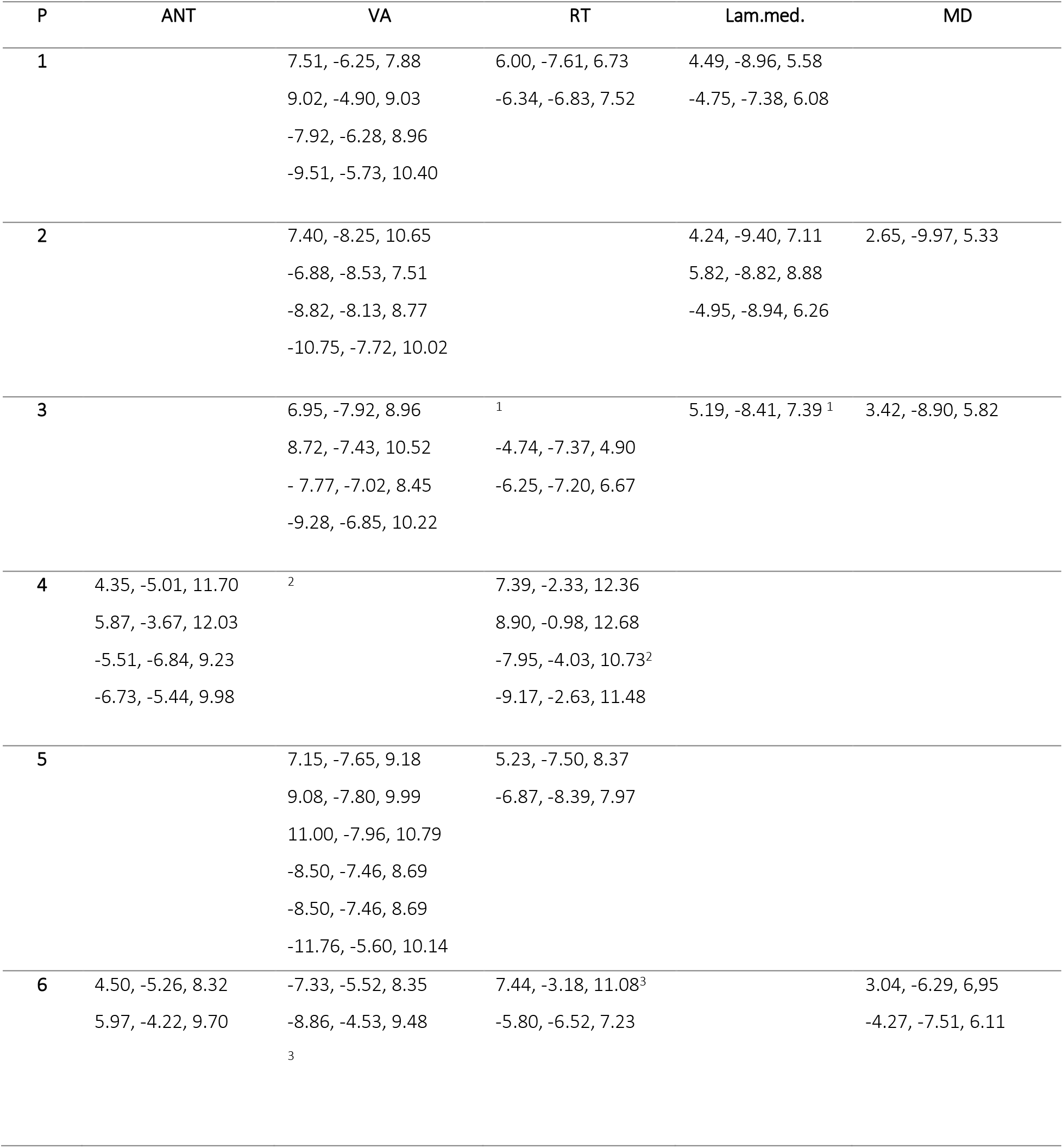
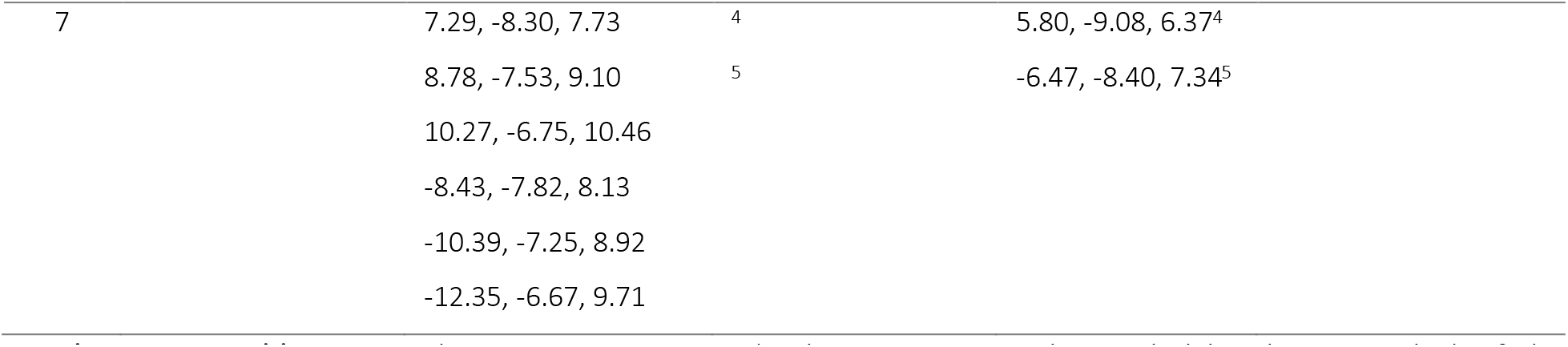
Coordinates in MNI-space (with respective nuclei guided by the Mai atlas) of the locations of electrode centers for all participants. Abbreviations: ANT: anterior nucleus of the thalamus; VA: ventral anterior nucleus; RT: Reticular nucleus; Lam.Med.: Lamina medialis; MD: mediodorsal nucleus Electrode locations at borders between nuclei are indicated by superscripts.

### Spindle characteristics per nucleus

We evaluated descriptive spindle measures when categorizing the data into their corresponding thalamic nuclei, selecting the three most common nuclei in our data. Note, that there was considerable uncertainty associated with the anatomical labeling. Three separate generalized linear mixed-effects models were constructed with density, duration, or amplitude as dependent variable, spindle type (slow vs. fast), thalamic nucleus (VA, ANT, RT) and their interaction as fixed effects, and participant as random intercept. The two-tailed significance level was set at α = 0.05. Mixed-effects models were computed using the Matlab function *fitglme*.

In the generalized linear model (GLM) analysis, there was a significant main effect of spindle type (F_1,60_ = 60.45; p < 0.001) on spindle density, but no main effect of nucleus (F_2,60_ = 1.48; p = 0.24) and no significant interaction of spindle type and nucleus (F_2,60_ = 2.29; p = 0.11) on spindle density. Post-hoc comparisons revealed that slow spindles (6.90 ± 2.88 sp/min) had significantly higher density (t(60) = 7.78; p < 0.001) than fast spindles (2.94 ± 1.52 sp/min). The intercept term was significant (F_1,60_ = 243.01; p < 0.001), pointing to inter-individual differences in spindle density.

When modelling spindle duration with nucleus, spindle type and their interaction as mixed effects, and participant as random intercept, there was no significant effect of spindle type (F_1,60_ = 4.01, p = 0.05), nucleus (F_2,60_ = 0.49, p = 0.62), or their interaction (F_2,60_ = 0.35; p = 0.71), but a significant intercept (F_1,60_ = 1033.5, p < 0.001). When modeling spindle amplitude, there was a significant main effect of spindle type (F_1,60_ = 6.80; p = 0.01), but no significant main effect of nucleus (F_2,60_ = 0.40, p = 0.67) and no interaction effect of spindle type and nucleus (F_2,60_ = 1.63; p = 0.20). Post-hoc comparisons revealed that slow spindles (9.96 ± 5.53 µV) had significantly higher amplitude (t(60) = 2.61; p = 0.01) than fast spindles (7.67 ± 5.85 µV) The intercept was significant (F_1,60_ = 36.47, p < 0.001), suggesting inter-individual differences in both spindle duration and amplitude.

**Supplementary figure 2.**
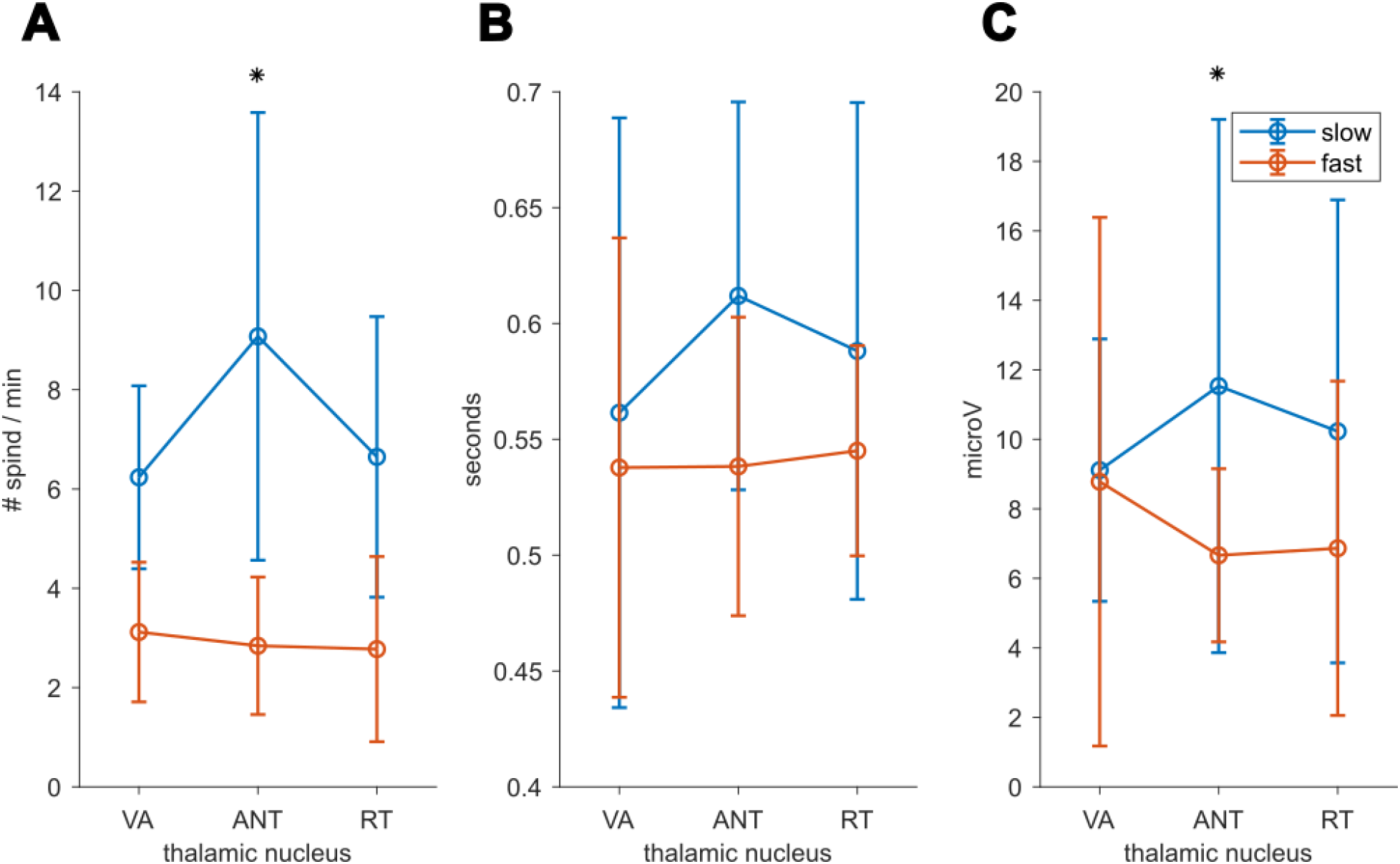
Average spindle characteristics per putative thalamic nucleus (see methods for limitations when estimating thalamic electrode locations). Values are shown for **A)** density; **B)** duration and **C)** amplitude. Asterisks mark significant results of spindle type at p < 0.05 in generalized linear mixed model. VA: ventral anterior nucleus of the thalamus; ANT: anterior nucleus of the thalamus; RT: Reticular nucleus of the thalamus.

### Simultaneous spindle onset in anterior thalamus and cortex

The previous analyses have shown that spindles tend to co-occur systematically. In order to understand this co-occurrence better, we investigated potential shifts in onset of thalamic and cortical spindles. Thalamic spindles might precede or follow cortical spindle activity. On the one hand, this would allow inferences on a causal relation of spindles between the two regions, but on the other hand this temporal shift would bias analyses of spatiotemporal spindle overlap toward channel pairs with a smaller temporal offset. The temporal shift was investigated by means of lag analysis. The lag was calculated as the time difference between spindle onsets in cortical relative to thalamic channels. Consequently, spindles with an earlier onset in the thalamus are indicated by a positive lag, and spindles with an earlier onset in cortical channels by a negative lag. Supplementary figure 3 depicts the median thalamocortical lag across participants and channels, for slow (A) and fast spindles (B), respectively. The distributions of thalamocortical lag for both slow (0.001 s ± 0.043) and fast (0.004 s ± 0.075) spindles were centered around 0. A one-sample t-test showed that both the slow (t(778) = 0.35; p = 0.72) and the fast (t(797) = 1.45; p = 0.15) spindles did not differ significantly from 0. This implies that spindle onset in thalamus and cortex was, on average, simultaneous.

**Supplementary figure 3.**
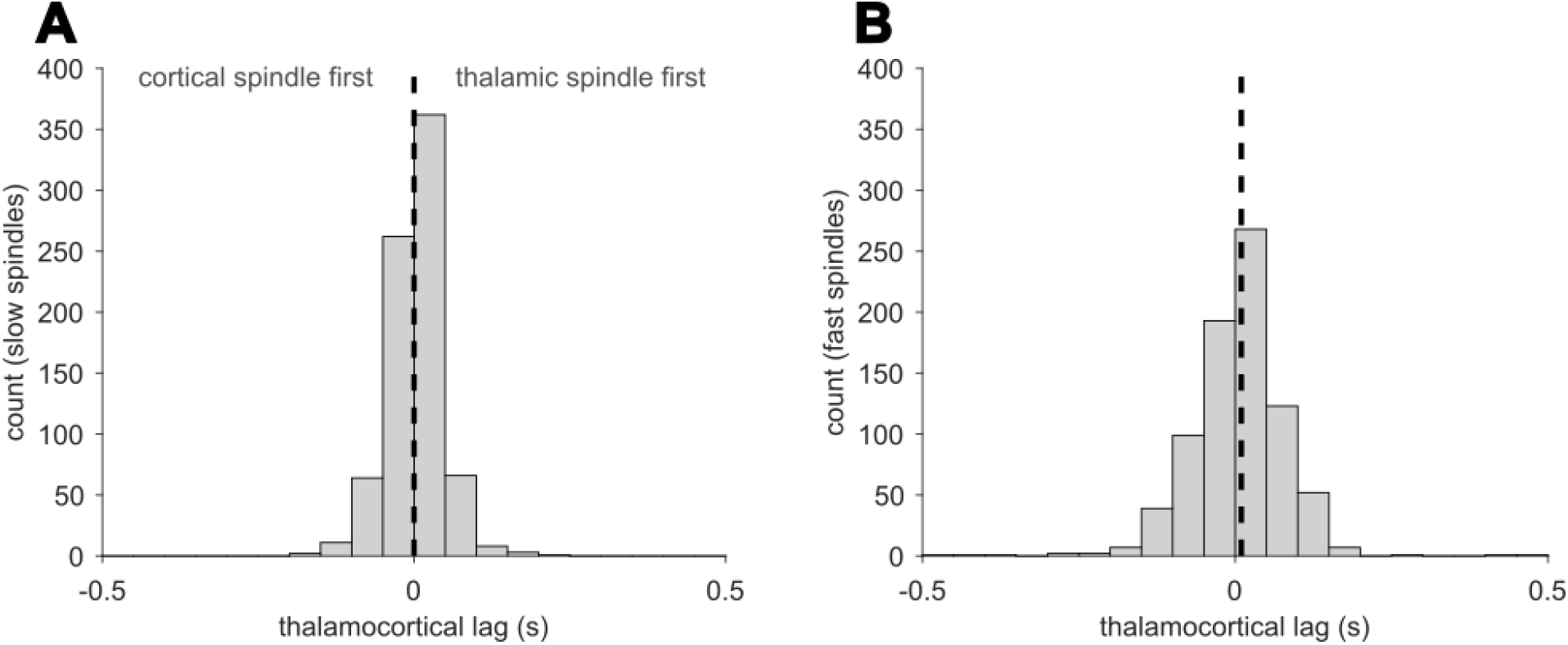
Histograms of thalamocortical lag of spindle onset across all thalamocortical channel pairs and participants. A positive lag indicates spindle onset in thalamus first, a negative lag in cortex first. **A)** slow spindles; **B)** fast spindles. Dashed line marks the median lag of cortical relative to thalamic spindle onset.

